# Multi-omics map revealed PPAR*α* activation protecting against myocardial ischemia-reperfusion injury by maintaining cardiac metabolic homeostasis

**DOI:** 10.1101/2023.08.17.551936

**Authors:** Yun Gao, Fei Huang, Fanwei Ruan, Dongwu Lai, Zhe Zhang, Yuan Zhang, Jun Zhu, Yuwen Lu, Liyin Shen, Jin He, Yan Liu, Guosheng Fu, Yang Zhu, Li Shen, Lenan Zhuang

## Abstract

Timely percutaneous coronary intervention is the most effective initial therapy for the acute myocardial infarction (MI). However, the mechanism in energy metabolism underlying time-dependent coronary reperfusion remains largely unknown. Here, we generated an integrative map of cardiac cells using bulk and single-nucleus RNA-seq combined with metabolomics profiling of hearts with reperfusion at distinct time points post MI in rat. We found early time reperfusion (ETR), but not late time reperfusion (LTR) reduced myocardial injury by maintaining cardiac energy homeostasis. PPARα was identified as a key regulator for maintaining fatty acid metabolism after MI/R injury. Importantly, pretreatment with FDA-approved PPARα agonist, fenofibrate, improved the transcriptional signatures, and ameliorated the function of the MI/R injured hearts, particularly in the ETR. Together, our data not only deciphered the protective effect of ETR by maintaining cardiac energy homeostasis, but also provided insights into the translational potential of PPARα activation in alleviating MI/R injury.

## Introduction

Acute myocardial infarction (AMI) caused by coronary artery occlusion remains as one of the major causes of morbidity and mortality worldwide, and presents a growing public health concern (*1*). The current paradigm for AMI therapy focuses on the repaid restoration of coronary artery blood flow to reestablish myocardial oxygen supply, known as reperfusion (*2*). In recent years, evidence showed that the early-reperfusion group had a lower incidence of arrhythmias and mortality rate, than the delayed-reperfusion and permanent-MI groups (*3, 4*). However, the underlying mechanism of myocardial energy metabolism remodeling in response to time-dependent coronary reperfusion remains unclear. Although substantial progress has been made in the acute therapy for myocardial infarction to help restore blood supply into the ischemic myocardium to reduce myocardial infarct size, reperfusion induces ischemia-reperfusion injury (IRI) to the myocardium (*5, 6*). The heart oxidizes a variety of substrates dependent on their availability in the circulation in response to external stimuli, allowing the heart to maintain pump function even in the case of metabolically abnormal to decrease energy deficiency switching substrate preference for ATP generation (from fatty acid to glucose utilization) (*7*). An ischemic heart has the ability to switch substrate preference from fatty acid to glucose oxidation with an increase in glucose uptake and glycolytic rates, frequently leading to decreased fatty acid oxidation and oxidative phosphorylation, and a compensatory increase in glycolysis, known as myocardial energy metabolism remodeling (*8-11*). In turn, the metabolic state of a cell regulates several cellular functions by activating intracellular signaling cascades. Thus, myocardial energy metabolism remodeling occupied with oxidative stress, reactive oxygen species (ROS) accumulation, calcium overload, and mitochondrial dysfunction contributes to cardiac myocardial IRI (*12-16*). Therefore, new approaches to limit myocardial IRI remain a significant unmet need for patients with AMI. Studies on normal, failed and partially recovered adult human hearts and pressure overload-induced pathological cardiac hypertrophy mouse heart have recently provided insights into cell-type-targeted intervention of heart diseases via single-cell and single-nucleus RNA sequencing (scRNA-seq and snRNA-seq, respectively) (*17-21*). Particularly, researchers create an integrative high-resolution map of human ischemic heart samples via scRNA-seq and snATAC-seq, together with spatial transcriptomics, yielding insights into changes in cardiac transcriptome and epigenome through the identification of distinct tissue structures of injury, repair and remodeling (*22, 23*). However, the regulation of myocardial energy homeostasis and cellular functions caused by cardiac remodeling after myocardial IRI remain unclear at the single-cell level. Furthermore, as the extent of myocardial IRI is largely dependent on time-dependent coronary reperfusion after acute MI, fast and efficient myocardial reperfusion remains the primary clinical goal. Researchers have found that substrate metabolism is regulated by the activation of specific transcription factors (TFs) to induce or suppress the expressions of important metabolic enzymes (*24, 25*). Among the activated TFs, PPARα plays a critical role in energy production and lipid as well as carbohydrate metabolism; it also regulates the expression of genes involved in peroxisomal and mitochondrial β-oxidation (*26, 27*). Peroxisome proliferator-activated receptor (PPAR), a superfamily of nuclear receptors, plays an important role in regulating metabolism process. Decreased fatty acid oxidation might be partially explained PPARα signaling suppression (*28, 29*). A recent study has found that PPARα activation prevents post ischemic contractile dysfunction in hypertrophied neonatal hearts by increasing the rates of cardiac fatty acid β-oxidation (*30*). PPAR activation requires co-activation with PPARγ co-activator 1α (PGC1α) or PGC1β, which are crucial regulators of mitochondrial biogenesis and cardiac mitochondrial function (*31-33*). Specifically, the PPARα-PGC1α axis plays an important role in regulating cardiac energy metabolism in healthy and diseased myocardium (*34*).

In this study, we created an integrative map of cardiac cells using bulk and single-nucleus RNA-seq combined with metabolomic profiling of reperfused hearts at distinct time points post infarction in rats. We found that ETR, but not LTR, reduced myocardial IRI by maintaining cardiac energy homeostasis. PPARα was identified as a key regulator in cardiomyocytes (CMs) for maintaining fatty acid metabolism after myocardial IRI. The metabolic state significantly changed the cellular function of NCMs in response to the myocardial IRI. Essentially, pretreatment with an FDA-approved PPARα agonist, fenofibrate, improved the transcriptomic profiles and the function of the IRI hearts, particularly in ETR. Together, these studies have revealed the relation to energy metabolism remodeling between the glucose and lipid metabolism in responding to IRI, and provided candidate therapeutic targets for future research and interventional opportunities to improve IRI treatment.

## Results

### Early time reperfusion reduced myocardial IRI by maintaining fatty acid metabolism

To investigate mechanisms underlying the myocardial IRI and the cardioprotective effects associated with reperfusion at various time points during myocardial IRI, the rats were subjected to left anterior descending coronary artery ligation for 1 h, 6 h (MI1h and MI6h) and 24 h (MI24h) along with Sham controls (Sham), and 1 h, 6 h ischemia followed by reperfusion (MI1h/R and MI6h/R, mimicking early time reperfusion (ETR) and late time reperfusion (LTR), respectively) followed by heart tissue collection 24 h post infarction to mimic the myocardial IRI in AMI patients (Fig. 1A). H&E staining and Masson’s trichrome staining revealed increased collagen deposition in LTR (MI6h/R) and prolonged myocardial ischemic hearts (MI24h) (Fig. 1B and fig. S1A, up and middle panels). Furthermore, the TUNEL assay showed that the MI1h and MI1h/R groups had less apoptotic cell than those in the MI6h/R and MI24h groups (Fig. 1B and fig. S1A, down panel). To determine the cardioprotective mechanisms underlying the different time points of reperfusion after myocardial infarction, RNA seq of the infarct zone from the Sham, MI1h, MI6h, MI1h/R, MI6h/R and MI24h groups was performed, and differentially expressed genes (DEGs) between the groups were identified (Fig1, C to E, fig. S1, B to E and table S1-2). Principal component analysis (PCA) revealed that MI surgery significantly changed the whole transcriptome compared with Sham. LTR exhibited similar gene expression profiles of the MI24h group but not ETR (Fig. 1C and fig. S1B). Interestingly, compared with LTR and the MI24h group, ETR was clustered closer with the Sham group. The heatmap demonstrated consistent results of all the groups by the expression levels of the DEGs between Sham and MI24h (fig. S1, C and D). Subsequently, we performed fuzzy c-means clustering of genes according to their expression profiles, which led to the identification of seven clusters (labeled as R-C1-C7; Fig. 1D and table S2). The R-C1 genes (n = 852) decreased with reperfusion delay and were associated with tricarboxylic acid cycle (TCA), mitochondrial transmembrane transport and fatty acid metabolic process. The R-C2 genes (n = 1670) were associated with immune response, autophagy, and response to oxidative stress. Genes related to DNA repair and B-cell immune response were detected in R-C3 (n = 1510). To determine the detailed differences between ETR and LTR, the DEGs between MI1h/R and MI6h/R were subjected to gene ontology (GO) analysis. Genes associated with mitochondrial gene expression and oxidative phosphorylation as well as mitochondrial respiratory chain complex I assembly were highly expressed in the MI1h/R group; however, these biological processes were inhibited in the MI6h/R group and further induced epithelium migration and wound healing (Fig. 1E). These results indicate that ETR (MI1h/R) is important for maintaining mitochondrial function to yield sufficient energy, which may account for protection against myocardial IRI of ETR. Quantitative reverse transcription polymerase chain reaction (qRT-PCR) was conducted to confirm the changes in the expression levels of representative genes. Genes involved in fatty acid metabolism (*Ppara*, *Ppargc1a* and *Cpt1b*) were reduced in the MI6h/R and MI24h groups. Consistently, myocardial IRI related genes (*Nppa*, *Nppb* and *Myh7*) were upregulated and cardiac function gene (*Cacna1c*) was downregulated in MI6h/R than in MI1h/R (Fig. 1F and fig. S1F). These results indicate that ETR is important for myocardial infarction in maintaining cardiac function and energy metabolism.

**Fig. 1.**
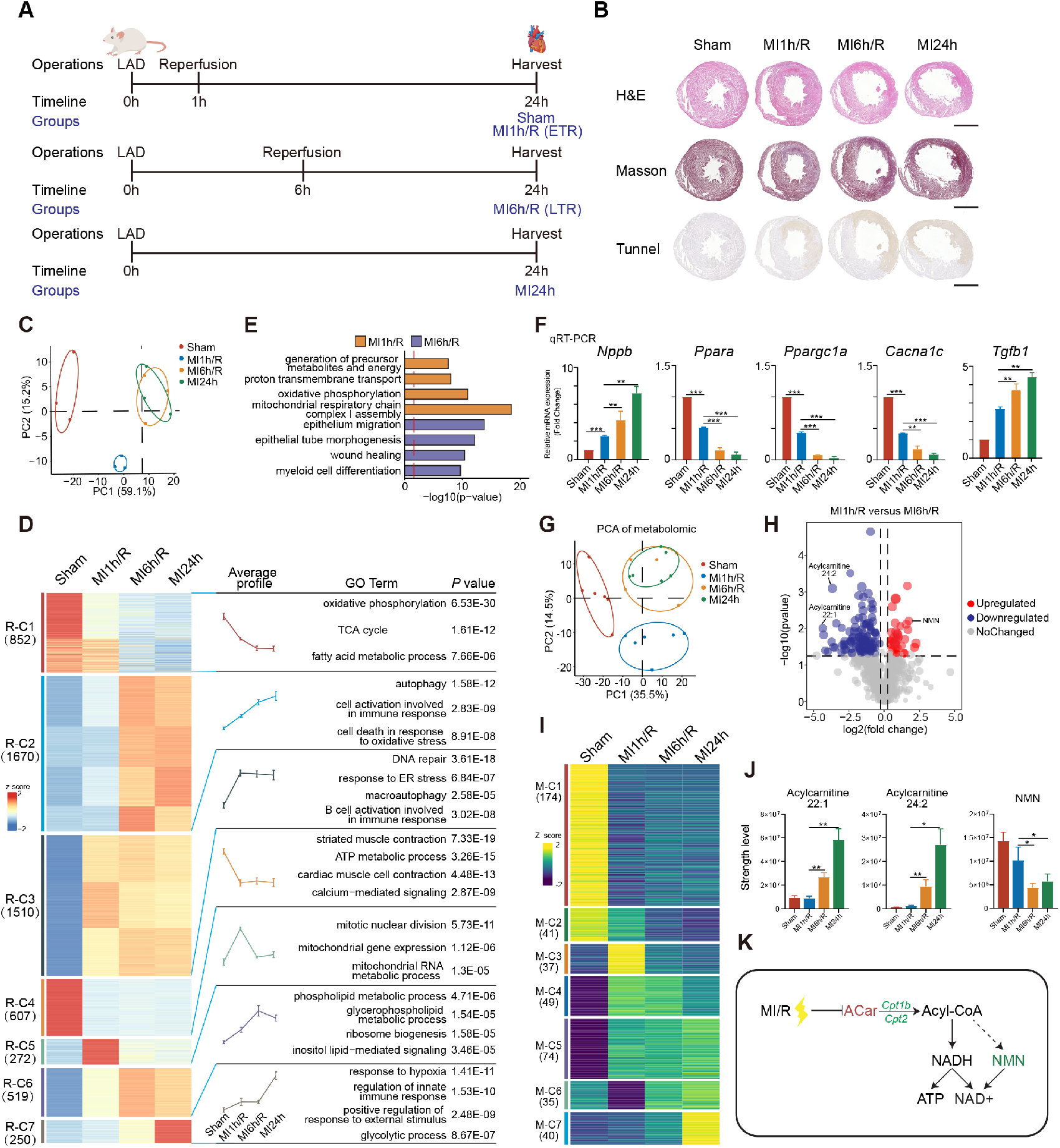
Early time reperfusion reduced myocardial IRI by maintaining fatty acid metabolism. **(A)** Schematic of study design. The rats were subjected to LAD coronary artery ligation for 1-hour, 6-hour ischemia followed by reperfusion for another 23 and 18 hours (MI1h/R and MI6h/R). **(B)** Histological analysis of Sham, MI and MI/R heart (up: H&E staining; middle: Masson trichrome staining; down: Tunnel assay). **(C)** PCA analysis based on the different periods of reperfusion. **(D)** Average expression levels and GO analysis of each clustered genes were shown by the heatmap and temporal profile of Sham, MI1h/R, MI6h/R and MI24h groups. **(E)** GO analysis of DEGs identified from the temporal myocardial ischemia/reperfusion groups. **(F)** The expression levels of key genes were measured by qRT-PCR. **(G)** PCA analysis of metabolomic data based on the different periods of reperfusion. **(H)** Volcano plots exhibiting DEMs identified from group MI6h and MI6h/R. **(I)** Average expression levels of each clustered metabolites were shown by the heatmap of Sham, MI1h/R, MI6h/R and MI24h groups. **(J)** Bar graphs showing relative abundance of the crucial metabolites among the four groups. **(K)** Schematic diagram of the metabolic flux highlighting the changes of metabolites between ETR and LTR. Green denotes down-regulated metabolites or gene (italics), red denotes up-regulated metabolites, and black denotes no change.

To further investigate the changes in whole cardiac metabolism, fresh rat hearts were collected for untargeted liquid chromatography-mass spectrometry (LC-MS) analyses. PCA of the metabolomic profile revealed that ETR grouped closely to Sham and clearly separated with LTR and that LTR grouped together with MI24h, which is consistent with the PCA results of transcriptome sequencing (Fig. 1G). Furthermore, 156 differentially expressed metabolites (DEMs) were identified from ETR and LTR (Fig. 1H). In the fuzzy c-means clustering, 7 clusters of metabolites (labelled as M-C1 to M-C7) were further identified according to the changes in their abundance (Fig. 1I and fig. S1G). Among the 156 DEMs, acylcarnitine (ACar), which played a key role in the β-oxidation of long-chain fatty acids through the inner mitochondrial membrane, had greater accumulation more in LTR and MI24h in the M-C7 cluster (Fig1, J and K). In addition, the NMN, which was involved in decreased myocardial injury, was significantly enriched in ETR than in LTR. Unexpectedly, decreased epigallocatechin was detected in LTR and MI24h (Fig. 1G), which was shown to be capable of decreasing mitochondrial injury and peroxidation in hypoxia-reperfusion-damaged CMs (*35*). Together, these results suggest that ETR maintained energy homeostasis alleviating myocardial IRI by upholding a suitable level of fatty acid oxidation.

### Single-nucleus transcriptomes revealed the cellular landscape of the cell composition of the IRI hearts

The size of mature CMs (>100 μm) were so large that unable to across the droplet-based platforms (maximum diameter is 40 μm), whereas increasing evidence shows that snRNA-seq revealed transcriptional signature of cardiac cells (*22, 36-38*). To investigate the cell-type-specific molecular mechanisms of temporal reperfusion-induced myocardial IRI, the heart infarct zones (Sham, MI1h/R, MI6h/R and MI24h) were subjected to single-nucleus transcriptome analysis. After the removal of low-quality cells, a UMI matrix of 22,234 cells and 17,003 genes was obtained (fig. S2a). To gain overall insights into the cellular heterogeneity of the IRI hearts, uniform manifold approximation and projection (UMAP) was employed to cluster all the nuclei (Fig. 2A and fig. S2B). The cell clusters were annotated by curated marker genes from the literature, ultimately, five major cell types were identified, which included namely, CMs, endothelial cells (ECs), fibroblasts (FBs), macrophages (MACs), and smooth muscle cells (SMCs) (Fig. 2B and table S3). Cell identities were validated by the expressions of cell-specific marker genes (Fig. 2, C to E and fig. S2C). Next, differences in the proportions of cardiac cells were analyzed. As expected, the Sham group had a larger proportion of CM, and the FBs were mainly derived from Sham and MI1h/R (Fig. 2, F and G). MI1h/R and MI6h/R occupied the high abundance of ECs. Interestingly, the proportion of MACs drastically increased in MI1h/R but decreased in MI6h/R and MI24h. Altogether, these results indicated that cardiac cell heterogeneity dynamically changed in temporal reperfusion induced myocardial IRI.

**Fig. 2.**
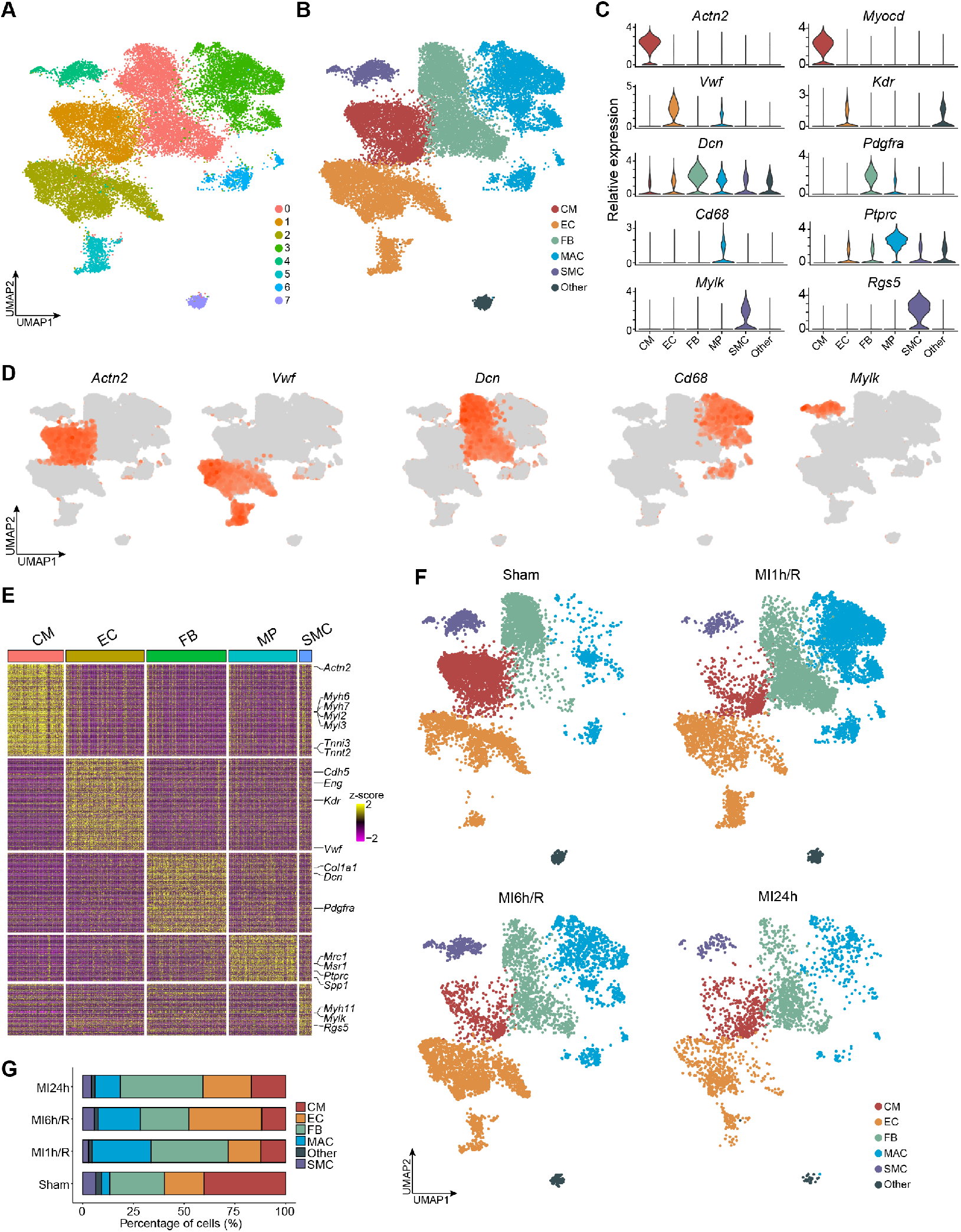
Single-nucleus transcriptomes revealed the cellular landscape of the cell composition of the IRI hearts. **(A-B)** UMAP plot of 22,234 single nuclei isolated from the Sham, MI1h/R, MI6h/R and MI24h groups. Cell types were identified by the expression of known marker genes. **(C)** Violin plots generated from the integrated dataset displaying characteristic marker genes of each identified cell population. **(D)** Feature plots showing expression of cell markers from CM, EC, FB, MAC and SMC. **(E)** Heatmap showing the molecular signature of each cell type. **(F)** UMAP plot showing cell contribution by the four group. **(G)** Fractional changes for each cell types across the Sham, MI1h/R, MI6h/R and MI24h groups.

### Early time reperfusion safeguarded PPAR*α*-mediated fatty acid metabolism in cardiomyocytes

CMs have been reported to play key roles in cardiac remodeling after myocardial infarction (*22*). To gain insights into the molecular heterogeneity of distinct CM states, all the CMs were aggregated into three CM subclusters based on transcriptional similarity using UMAP (Fig. 3A and fig. S3, A to C). A total of 1461 genes were upregulated in CM1, whereas 1546 genes were highly expressed in CM3 (fig. S3D and table S4). Interestingly, 82% of the CM2-upregulated genes overlapped with the CM1-upregulated ones (1,138 of 1,385), indicating that the CM2 subclusters was similar to CM1 but not to CM3 (fig. S3D and table S5). Among these three CM subclusters, the representative genes exhibiting significant changes in centrality were identified (Fig. 3B). GO analysis revealed that 1,461 CM1 upregulated genes were associated with heart contraction and lipid-related energy metabolism, whereas 1,546 CM3-upregulated genes were associated with physiological stress response and immune response (fig. S3E). Comparison between metabolic pathway expression and the CM types enabled us to link information on cell metabolism to the function of the three CM subclusters. The results indicated that CM1 and CM2 showed significant similarities in terms of increased lipid oxidation, TCA, heart contraction, and decreased glycolysis signaling expression, whereas CM3 had opposite signaling activities (Fig. 3C). Gene set variation analysis (GSVA) showed that heart contraction and lipid metabolism pathways were activated in CM1 and then in CM2 but inactivated in CM3. However, glycolysis, hypoxia, and ECM pathways significantly increased in CM3 (Fig. 3D and fig. S3F). Furthermore, CM1 and CM2 exhibited high expression levels of lipid metabolism-related genes (*Ppara*, *Ppargc1a* and *Cpt1b*) and heart function genes (*Myh6* and *Mybpc3*), however, *Eno1*, encoding glycolytic enzyme, pathological cardiac hypertrophy gene *Nppa*, and fibrosis-related gene *Col1a1* were highly expressed in CM3 (Fig. 3, E and F and fig. S3G). These results indicated that the three CM subclusters exhibited different characteristics and cell states: 1) CM1, “healthy” CMs; 2) CM2, “subhealthy” CMs; 3) CM3, “injured” CMs. This is consistent with the results of the cellular composition comparison between the sample groups, which indicated that the CM1 and CM2 populations were mainly derived from the Sham and MI1h/R groups, respectively. CM3 was attributed to MI6h/R and MI24h (Fig. 3G and fig. S3H). To define the regulatory networks directing the three different CM states, we conducted TF binding prediction analysis around subclusters-specific gene promoters (1,138 shared upregulated genes in the CM1/2 and 949 genes in CM3). The PPARα motif was significantly enriched in the CM1/2-upregulated gene promoters, whereas the Klf1/2 motif was enriched in the promoters of the 1546 CM3-upregulated genes (Fig. 3H). In addition, TF-target gene regulatory network analysis of the three CM subclusters was employed to identify major TFs. In this analysis, PPARα played a major role in the regulatory module associated with healthy and subhealthy CMs (fig. S3I). Furthermore, the target genes of *Ppara*, including *Cacna1c*, *Ppargc1a*, and *Cpt1b*, were highly expressed in CM1 and CM2, whereas the central regulator for the CM3-enriched genes was *Zeb2*, a key regulator of epithelial-mesenchymal transition. PPARα and its target genes were defined as the “PPARA signature”. CM1 and CM2 had high expression of the PPARA signature, whereas CM3 had a much lower expression (fig. S3J). Similar results can be obtained from reanalysis of the single cell transcriptome data of human ischemic hearts, which the CMs were clustered into nonstressed vCM1, prestressed vCM2, and stressed vCM3(*22*). vCM1 and vCM2 had high expression of the PPARA signature, whereas the stressed vCM3 had a lower expression (fig. S3K). Immunofluorescence staining was performed to confirm the differential expression level of NPPA in each group. NPPA significantly increased in the infarcted area of the MI6h/R and MI24h groups (Fig. 3I). The expression level of PGC1α in the infarcted area was also evaluated, and PGC1α was found to be significantly higher in Sham and MI1h/R than in MI6h/R and MI24h (Fig. 3J). To infer the future fate of CMs, RNA velocity analysis on the UMAP embeddings was conducted in the aforementioned four groups(*39, 40*). Interestingly, a strong directional flow from CM2 to CM1 was only observed in the MI1h/R group, indicating that the CM2 from the ETR group has a transition potential to the healthy state (Fig. 3K). To further elucidate the mechanism of the directional flow from CM2 to CM1, cells in transiting state were extracted to evaluate transcriptional profile (Fig. 3L and fig. S3, L and M). The highly expressed genes in this state were enriched in heart contraction and response to mechanical stimulus (fig. S3L). Subsequently, some calcium ion channel regulators (*Ryr2*, *Cacna1c*) and heart contraction related genes (*Myom2*, *Tnni3k*) were found to exhibit significant transcriptional dynamics (Fig. 3L and fig. S3M). Interestingly, these transiting driver genes were all PPARα target genes identified in the previous analysis. These results indicated that the inferred directionality of CM2 to CM1 is mainly governed by PPARα and its target genes. Overall, these findings indicated a significant change in the TF regulatory network and CM states along the temporal reperfusion-induced metabolic remodeling. The nuclear receptor PPARα may play a key role in mediating fatty acid oxidation remodeling and calcium ion homeostasis in response to myocardial IRI.

**Fig. 3.**
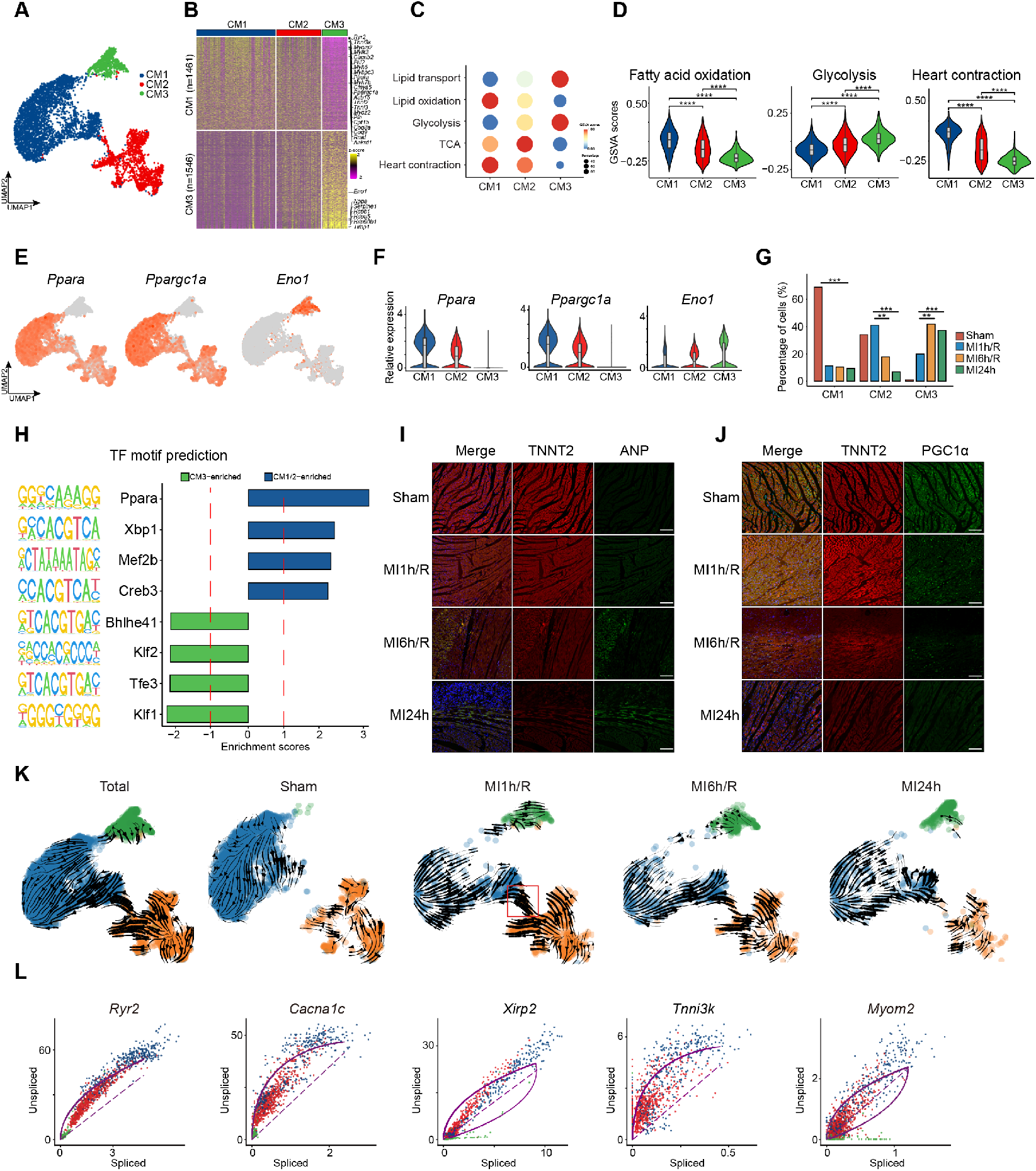
Early time reperfusion safeguarded PPAR*α*-mediated fatty acid metabolism in cardiomyocytes. **(A)** UMAP projection showing 5,239 CMs isolated from infarct zone of rat hearts at different times of reperfusion. Cells are colored by subclusters. **(B)** Heat map showing enriched genes identified by comparing CM1 and CM3. **(C)** Dot plot of expression scores for metabolic process of the 3 CM subclusters. **(D)** GSVA analysis of the important signaling pathways. **(E-F)** Violin plots **(E)** and Feature plots **(F)** showing expression of *Ppara*, *Ppargc1a* and *Eno1*. **(G)** Statistical analysis of the subclusters CM populations. **(H)** The corresponding enriched promoter-binding motifs of transcript factors identified from 3 CM subclusters. **(I)** Immunostaining of TNNT2 and NPPA was used to determine the extent of myocardial damage. **(J)** Immunostaining of TNNT2 and PGC1α was used to determine the location of crucial genes. **(K)** The UMAP projection shows the landscape of the process of in total and four groups of Sham MI1h/R, MI6h/R and MI24h. **(L)** The dynamical model allows to systematically identify putative driver genes.

### Heart function and heart disease signatures were identified to assess the cardiac states in human patients

According to the aforementioned analysis, we have found that the three CMs subclusters were in various levels of damage: healthy, subhealthy and injured. These CMs significantly differed in glucose and lipid metabolism, which was favorable for setting gene expression signatures to access the myocardial states in other experiments. To select genes for the efficient evaluation of myocardial states, one gene set composed of 140 genes (overlap between the 1138 CM1/2-shared upregulated genes and bulk RNA-seq-Sham high genes) was defined as the “heart function” signature; another 149 genes (overlap between the 949 CM3-upregulated genes and bulk RNA-seq-MI24h-high genes) were defined as the “heart disease” signature (Fig. 4, A and B). The heart function genes were enriched in heart contraction, calcium-mediated signaling, and fatty acid oxidation (fig. S4A). Contrarily, the heart disease signature genes were associated with immune response and wound healing. CM1 and CM2 had significantly higher expression of the heart function signature than CM3; in addition, and the representative heart function gene *Ryr2* was highly expressed in these CMs (Fig. 4, C and D). A reduction in the heart disease signature in CM1 and CM2 compared with CM3 was observed, and the representative heart disease gene *Anp* was highly expressed in CM3 (Fig. 4, E and F). To decipher the transcriptomic dynamics during transition from healthy toward injured CMs, a trajectory was reconstructed through pseudo-temporal ordering of three CM subclusters using Monocle2 (*41*). In lineage 1, the root state, CM1 of Sham exhibited a skewed cell distribution toward CM2 of MI1h/R, subsequently reached the end point, CM3 composing of MI6h/R, indicating that ETR, not LTR, is similar to healthy CMs. In lineage 2, a simple healthy-to-injured trajectory was observed, which started with CM1 of Sham while and ended with CM3 of MI24h (Fig. 4, G and H and fig. S4B). The calculated pseudo-time of the trajectory path exhibited LTR and MI24h-enriched CM3 toward later pseudo-time, which was in line with the development of myocyte injury (Fig. 4H and fig. S4B). Lipid metabolism associated genes (*Ppara*, *Ppargc1a,* and *Cpt1b*) and heart function-related genes (*Myh6*, *Mybpc3*) decreased along with the pseudo-time (Fig. 4I and fig. S4C). Furthermore, glycolytic enzyme *Eno1*, myocardial damage landmark genes *Anp* and *Ankrd1*, and fibrosis-related genes (*Spp1*, *Col1a1,* and *Tgfb1*) showed a sharp increment with the pseudo-time (Fig. 4I and fig. S4D). In agreement with the healthy-to-injured state transition of CM, the heart function signature had the similar expression trend (Fig. 4J). The myocardial damage of bulk transcriptomic data of heart tissues using the aforementioned two signatures was assessed further, and the results indicated that 1 h myocardial ischemia caused only minimal damage followed by ETR (Fig. 4K and fig. S4E). However, short-time myocardial ischemia and late reperfusion caused equivalent severity of myocardial IRI. To validate the confidences of heart function and heart disease signatures, the two signatures were evaluated using the data of human ischemic heart samples (*22*), and high agreement and correlation in terms of vCM cell states were observed (Fig. 4L and fig. S4, F and G). Of note, the expression of the two signatures was evaluated in the spatial transcriptomic profile of human ischemic heart samples. The two signatures clearly distinguished the 3 different CMs enriched zones of the human samples. Heart function highly expressed in the non-stressed vCM1 and pre-stressed vCM2 located regions, whereas heart disease showed high expression in stressed vCM3 located region (Fig. 4M and fig. S4, H and I). Together, the heart function and heart disease signatures, identified from our bulk and single-nucleus RNA-seq data, served as a good standard to evaluate CM states and tissue injury regions.

**Fig. 4.**
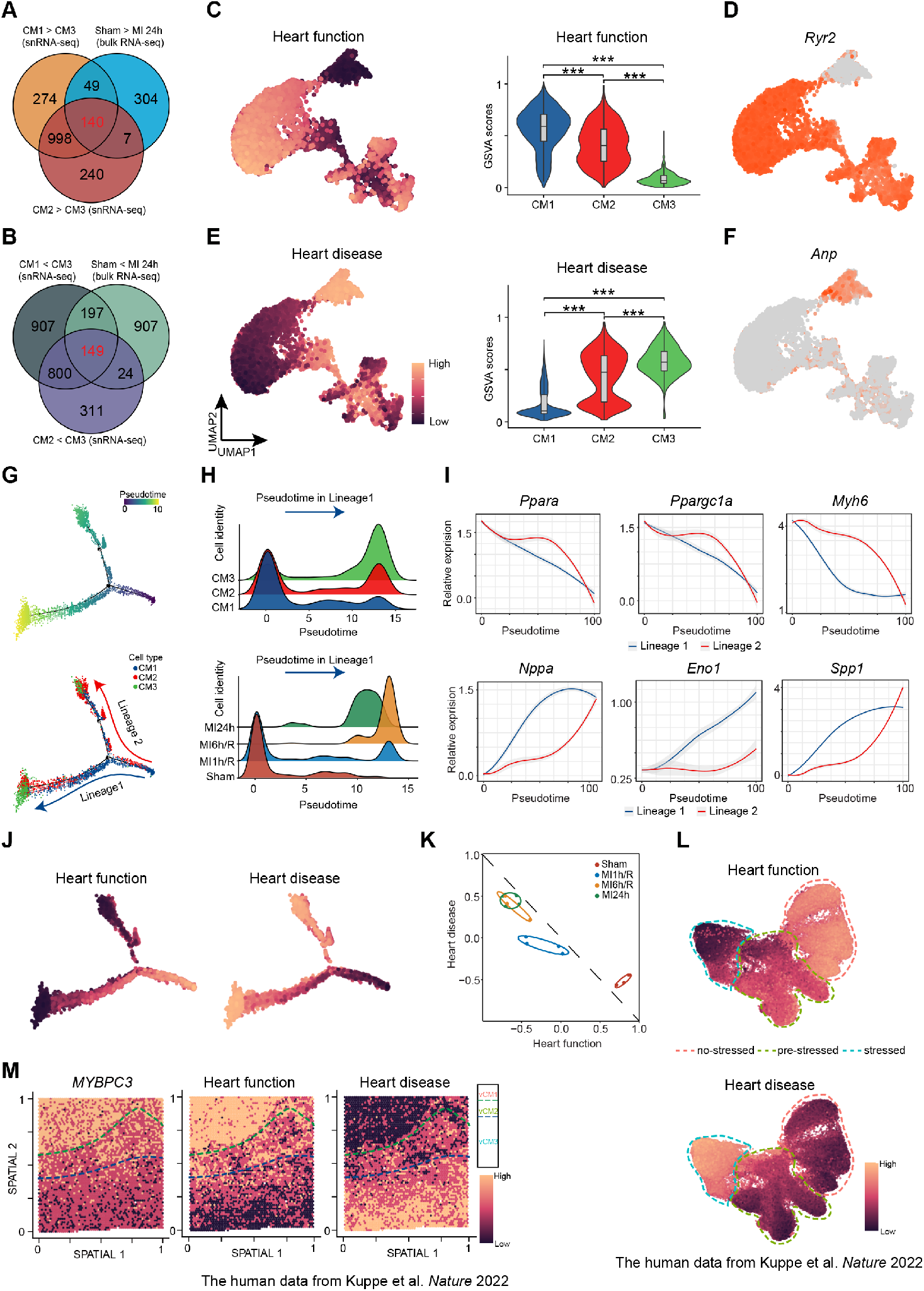
Heart function and heart disease signatures were identified to assess the cardiac states in human patients. **(A)** The number of the overlapping genes specifically upregulated in CM1/2 (compared with CM3) and Sham group (compared with MI24h) to identify heart function genes. **(B)** The number of the overlapping genes specifically upregulated in CM3 (compared with CM1/2) and MI24h group (compared with Sham) to identify heart disease genes. **(C-D)** Feature plot and violin plots of the expression level of heart function **(C)** and feature plot showing the expression level of representative heart function gene in CMs **(D)**. **(E-F)** Feature plot and violin plots of the expression level of heart disease **(E)** and feature plot showing the expression level of representative heart disease gene in CMs **(F)**. **(G)** Diffusion pseudo-time analysis of the CM lineage. The order in the diffusion pseudo-time (the up panel), CM subclusters (the down panel). **(H)** Density plots for cell density of the 3 CM subclusters (the up panel) and the four groups (the down panel) along with the pseudo-time lineage 1. **(I)** Expression for the representative heart function (the up panel) and heart disease genes (the down panel) over pseudo-time. **(J)** Feature plot showing the expression of heart function (the up panel) and heart disease signatures (the down panel) representative genes along with the course of pseudo-time. **(K)** The analysis of heart function and heart disease signatures based on the bulk tissues transcriptomics datasets. **(L)** The identification of heart function and heart disease signatures in the published human myocardial infarction heart single-cell sequencing datasets. **(M)** The identification of heart function and heart disease signatures in the published human myocardial infarction heart spatial transcriptomics datasets. RZ, remote zone; BZ, border zone; IZ, ischemic zone.

### Metabolic remodeling affected the function of cardiac fibroblasts and endothelial cells

Unsupervised clustering of FBs from all samples identified three FB subclusters (FB1-3), and differed in their abundance and transcriptional patterns (fig. S5, A to C). The glucose and lipid metabolisms across the FB subclusters were analyzed. ETR- and LTR-enriched FB2 induced a strong lipid metabolism accompanied by glycolysis, which further stimulated FB migration and inflammatory response (Fig. 5A). Nevertheless, FB3 exhibited high expression of glycolytic genes and the highest ECM score with the high expression of *Postn* (Fig. 5, A to C). Essentially, the energy metabolism remodeling of FBs caused different functional collagens. FB1 mainly expressed network-forming collagens to maintain FB cytoskeleton. FB3 had strong expression of fibrillar and fibril associated collagens with interrupted triple helices after myocardial IRI (Fig. 5D). Furthermore, the potential differentiation path of the three FB subclusters was identified to calculate trajectories and significant differences were observed in the pseudo-time distributions with unique expression profiles of *Postn*, *Top2a, Vwf,* and *Tgfb1* (Fig. 5, E and F and fig. S5D). All the results indicated that energy metabolism remodeling induced the shift of the FB populations toward an ECM-expressing profile in response to myocardial IRI.

**Fig. 5.**
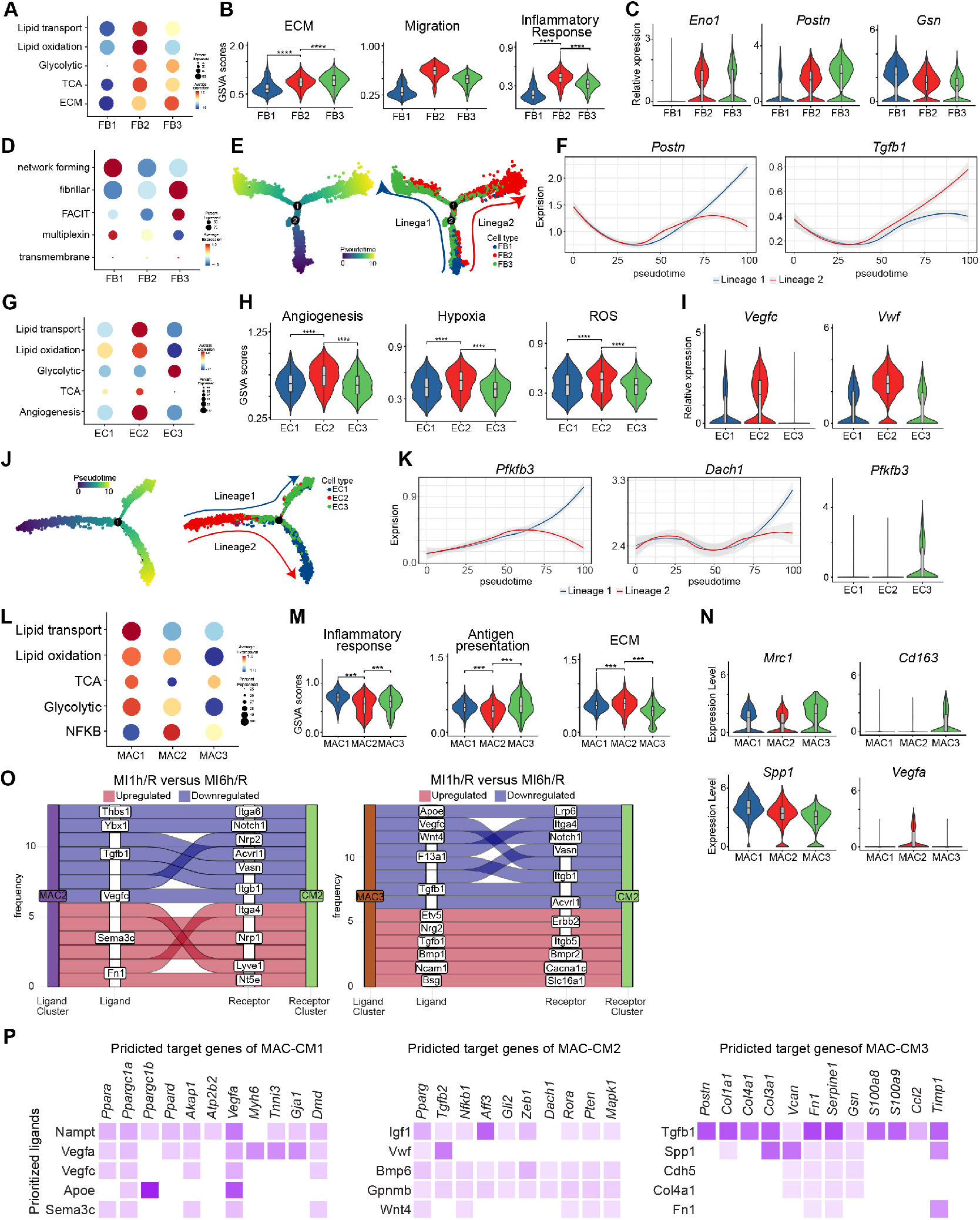
Metabolic remodeling affected the function of FBs, ECs and MACs as well as the cell-cell interactions of cardiac cells. **(A)** Dot plot of expression scores for metabolic process of the 3 FB subclusters. **(B)** GSVA analysis of the important signaling pathways. **(C)** Violin plots of crucial genes. **(D)** Dot plot of expression scores for collagen subtypes per FB subclusters. Dot size refers to proportion of cells expressing the gene set per cluster. **(E)** Diffusion pseudo-time analysis of the FB lineage. **(F)** Expression for the representative genes over pseudo-time. **(G)** Dot plot of expression scores for metabolic process. **(H)** GSVA analysis of the important signaling pathways of the 5 EC subclusters. **(I)** Violin plots of crucial genes. **(J)** Diffusion pseudo-time analysis of the EC lineage. **(K)** Lipid metabolism genes-related gene expression for *Acsl5* and *Dach1* over pseudo-time separated by four groups. **(L)** Dot plot of expression scores for metabolic process of the 3 MAC subclusters. **(M)** GSVA analysis of the important signaling pathways. **(N)** Violin plots of crucial genes. **(O)** Sankey plots of ligand-receptor interactions for MAC2/3 and CM2 identified from MI1h/R versus MI1h/R. These ligand-receptor pairs are selected by absolute value of the difference in LRscore. **(P)** The heatmap of select ligands expressed by MAC-CM1 (rows of the left panel), MAC-CM2 (rows of the middle panel) and MAC-CM3 (rows of the right panel) to regulate the genes which are differentially expressed by the CM1 (columns of the left panel), CM2 (columns of the middle panel) and CM3 (columns of the right panel). Well-established ligand-target gene interactions shown with a darker shade of purple.

The cardiac ECs also played a crucial role in promoting CM proliferation and maturation via paracrine signaling (*42, 43*). Unsupervised clustering of ECs from all samples identified five EC subclusters (EC1-5), and differed in their abundance and transcriptional patterns (fig. S5, E to G). GSVA revealed that genes associated with lipid transport and lipid oxidation were significantly upregulated in EC2, which were involved in angiogenesis with the high expression of angiogenic genes *Vegfc* and *Vwf* (Fig. 5, G to I). However, the lipid metabolism was downregulated and the glycolysis process upregulated in EC3 highly expressed glycolytic gene *Pfkfb3* (Fig. 5, G to I). Pseudo-time analysis revealed two lineages with different major cell compositions along with the pseudo-temporal ordering of ECs (Fig. 5J). EC2 was the least differentiated cluster, whereas EC3 was the most differentiated one. *Dach1*and *Pdkdb3* exhibited high expressions in lineage1 (Fig. 5, J and K and fig. S5H). Together, these findings highlight the different glucose and lipid metabolism induced multiple functional EC subclusters and their potential roles in angiogenesis.

### The heterogeneity and cell-cell interactions of macrophages contributed to remodeling of cardiac cells

To understand the heterogeneity of macrophages during myocardial IRI, three macrophage subclusters were identified, and differed in their abundance and transcriptional patterns (fig. S5, I to K). GSVA indicated that MAC1 exhibited high lipid and glucose metabolism, which was associated with increased inflammatory response (Fig. 5, L and M). MAC2 had a high expression of the glycolysis process, whereas the MAC3 had high expression of genes associated with TCA and antigen presentation (Fig. 5, L and M). MAC1 was involved in wound healing and phagocytosis with a high expression of *Spp1*, whereas MAC2 was associated with cell migration with high expression of *Vegfa*. MAC3 had high expressions of resident MAC markers *Cd163, Lyve1,* and *Mrc1*, which were associated with immune response, suggesting that MAC3 had anti-inflammatory effects (Fig. 5N). The metabolic states of macrophages not only provided sufficient energy but also reprogramed its phenotypical and functional alterations.

To investigate the role of different subclusters of macrophages during myocardial IRI and remodeling by cellular communication, receptor-ligand interaction analysis of CM/noncardiomyocyte (NCM)-MAC was calculated. Interestingly, distinct changes in cellular crosstalk were observed between ETR and LTR. Receptor-ligand pairs Ncam1-Cacna1c and Nrg2-Erbb2 increased, whereas Wnt4-Notch1 and F13a1-Itgb1 decreased among the interaction of MAC3-CM2 in ETR versus LTR (Fig. 5O). Of note, ETR exhibited increased Vegfc-Lyve1 and Fn1-Nt5e of MAC2-CM2, whereas LTR showed increased Vegfc-Itgb1 and Ybx1-Notch1 signal. These results indicated that the MAC-CM interaction was a key event and that the energy metabolism homeostasis of MI1h/R favored anti-inflammatory response during myocardial IRI. Furthermore, target gene co-expression analysis was conducted using the NicheNetR algorithm to determine possible target genes(*44*). Nampt, Vegfa and Sema3c were key regulatory ligands in MAC to interact with receptor in CM1 and Vwf, Gpnmb, and Wnt4 in CM2, whereas ECM-related genes Tgfb1 and Spp1 were key regulatory ligands in CM3 (Fig. 5P). *Ppara*, *Ppargc1a*, *Ppargc1b* and *Ppard* were highly expressed in the “receiver” CM1 and CM2, which was predicted to be caused by the presence of Nampt, Apoe, Igf1 and Bmp6 from MACs, but not in CM3. Meanwhile, cellular crosstalk analysis between MACs and FB, EC was conducted. Tgfb1 and Hmgb2 were important ligands affecting the expressions of *Eln*, *Top2a* and *Mki67*, which enhanced the activation of cardiac FBs contributing to fibrosis (fig. S5, L and M). Furthermore, enhanced Vegfc-Nampt, downregulated Vegfa-Fgf1 among the interaction of MAC-EC were found in ETR versus LTR samples. The angiogenesis marker Vegfa was the important ligand from MAC2 that influenced the expression of the downstream genes (*Hgf* and *Nos3*) in EC2 to promote angiogenesis (fig. S5, N and O). These results indicated that the metabolic state significantly reprogramed the cellular heterogeneity and cell-cell interactions of MACs in response to myocardial IRI.

### Pretreatment with fenofibrate ameliorated myocardial IRI

Previous analysis demonstrated the crucial role of PPARα in maintaining fatty acid metabolism in response to myocardial IRI. Fenofibrate was used to test the protective effect of PPARα activation on myocardial IRI. Rats received either fenofibrate (100 mg kg^-1^; intraperitoneal injection) or PBS 2h before MI surgery (Fig. 6A). Transcriptome analysis was conducted, and the results were compared with our previous data. Both PCA and heatmap analysis revealed that fenofibrate treatment changed the transcriptional patterns of the IRI hearts (fig. S6, A and B). Interestingly, the heart function and heart disease signatures well coherentized the groups, and the fenofibrate treatment upregulated the expression of heart function genes and downregulated the heart disease genes (Fig. 6B and fig. S6, C to E). Importantly, fenofibrate significantly upregulated genes associated with heart contraction, fatty acid oxidation and TCA pathways (Fig. 6C). Highly expressed genes identified in FBs (*Tgfb1* and *Ccn2*), ECs (*Pdgfa* and *Cxcr2*), and MACs (*S100a8* and *S100a9*) also decreased after fenofibrate treatment. Western blot assays revealed that fenofibrate induced PPARα protein expression, particularly in ETR, and that PGC1α protein was higher in response to myocardial IRI in fenofibrate-treated hearts, particularly in ETR, the Cpt1b exhibited similar results (Fig. 6D). These results indicated that fenofibrate significantly upregulated PPARα expression to maintain energy metabolism allowing further improvements, which appeared to be more effective with ETR. To further determine whether the fenofibrate-activated PPARα pathway would improve myocardial outcome after myocardial IRI, rats that received either fenofibrate or PBS were followed up weekly with echocardiography until 4 weeks after the myocardial IRI (fig. S6F). The fenofibrate-treated rats showed less infarct size, thicker infarcted walls, and fibrosis than those without fenofibrate treatment (Fig. 6E, fig. S6G). Importantly, echocardiography revealed that fenofibrate-treated rats had significantly higher left ventricular ejection fraction and fractional shortening than the PBS-treated rats, particularly in MI1h/R 7dpi (7 days post infarction) and 28dpi (28 days post infarction) (Fig. 6, F to I, fig. S6, H and I). Altogether, these results indicated that PPARα pathway activation via fenofibrate improved the transcriptional signatures and function of the IRI hearts in vivo.

**Fig. 6.**
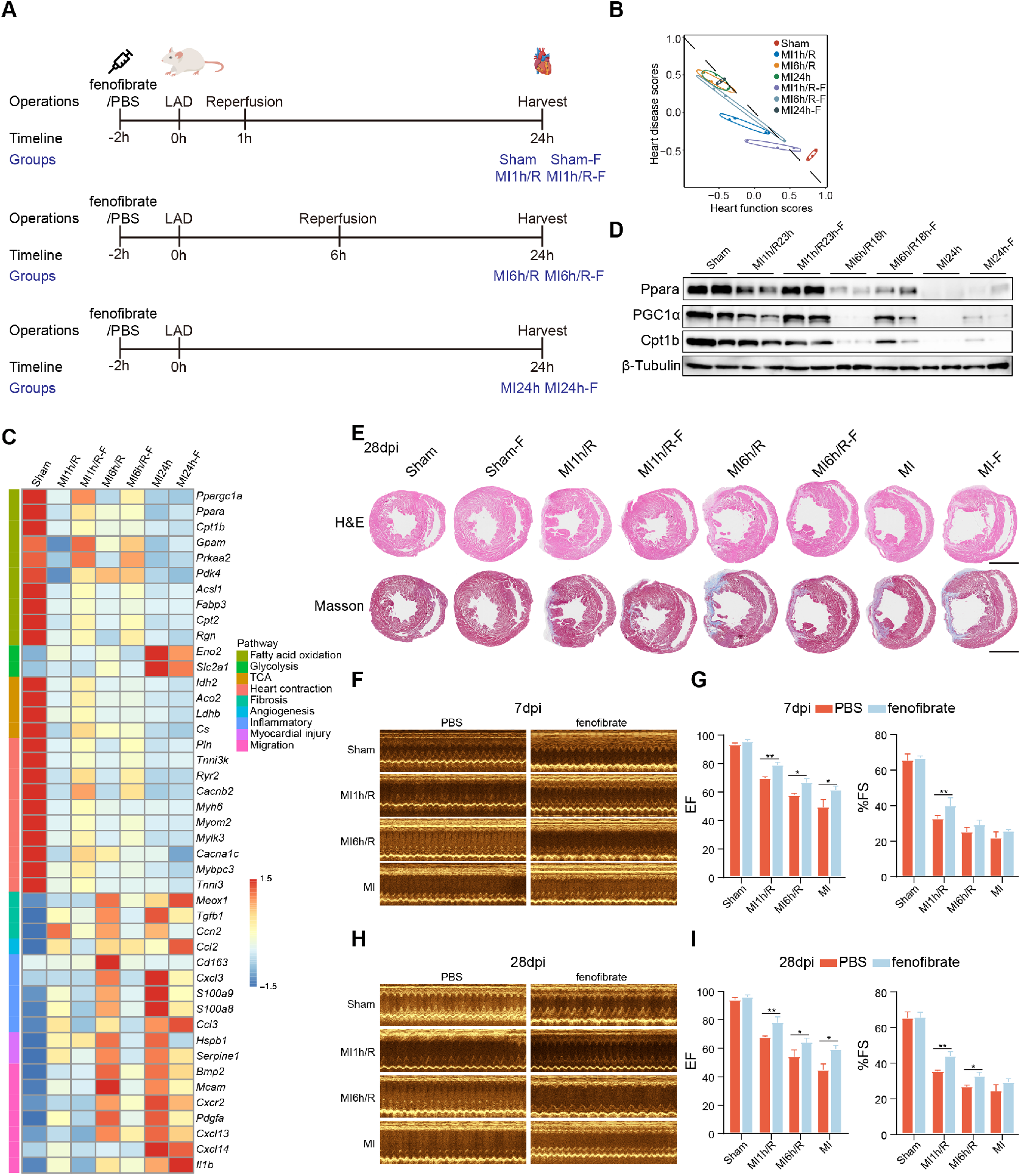
Pretreatment with fenofibrate ameliorated myocardial IRI. **(A)** Schematic of study design and the rats were treated with PPARα agonist fenofibrate 2 hours before LAD. **(B)** The analysis of heart function and heart disease signatures based on the bulk tissues transcriptomics datasets. **(C)** Heatmap showed crucial genes of important pathways with PPARα agonist fenofibrate. **(D)** Western blotting of the crucial protein in the infarction zone. **(E)** Histological analysis of Sham, MI and MI/R heart(Up: H&E staining; Down: Masson trichrome staining). 28dpi: 28 days post infarction. **(F)** M-mode echocardiography images of rat 7 days post infarction (7dpi). Results are representative of five different samples. **(G)** Each bar plot represents the left ventricular ejection fraction (left) and fractional shortening (right) of the different groups 7 days post infarction. **(H)** M-mode echocardiography images of rat 28 days post infarction. Results are representative of five different samples. **(I)** Each bar plot represents the left ventricular ejection fraction (left) and fractional shortening (right) of the different groups 28 days post infarction.

## Discussion

In this study, an integrative map of cardiac cells was generated using bulk and single-nucleus RNA-seq combined with metabolomic profiling of reperfused hearts at different time points post infarction in rats, which provided insights into the cellular composition and interaction networks of the heart with various reperfusion time points. Compared with the no-reperfusion control, ETR, but not LTR, reduced myocardial IRI by maintaining cardiac energy homeostasis. PPARα was identified as a key regulator for maintaining fatty acid metabolism. Pretreatment with fenofibrate, an FDA-approved PPARα agonist, improved the transcriptional signatures and function of the IRI hearts, particularly in ETR.

Previous studies have proposed several molecular mechanisms including energy metabolism remodeling, ROS release, calcium overloading and programmed cell death activation for the pathological alterations in myocardial IRI (*45-49*). However, the mechanism in energy metabolism remodeling underlying the time-dependent coronary reperfusion therapy of IRI remains largely unknown. In this study, an integrated multi-omics analysis combining transcriptomics and metabolomics was conducted. We found that *Ppara*, *Ppargc1a,* and *Cpt1b*, which were fund to play important role in fatty acid oxidation and oxidative phosphorylation, were significantly downregulated in LTR and MI24h, consistent with the accumulation of ACar and decrease in NMN in LTR and MI24h. These results indicated that myocardial IRI was induced by the dysregulated energy metabolism in LTR and MI24h, whereas ETR still maintained steady energy metabolism similar to Sham. The multi-omics analysis revealed that the major contributor to the myocardial IRI was fatty acid oxidation blockage (ACar accumulation). ETR alleviated myocardial IRI by consuming fatty acid to generate sufficient energy, which is consistent with the finding of a previous study indicating that the ability to increase fatty acid oxidation during post-ischemic reperfusion is important for myocardial recovery post ischemia (*30*). Heart function and heart disease signatures were proposed based on the bulk RNA-seq and snRNA-seq, which were employed to precisely and comprehensively evaluate myocardial injury. The heart function genes were enriched in terms of fatty acid oxidation, heart contraction, and calcium-mediated signaling, which further demonstrated that energy metabolism remodeling occupied an important portion of myocardial injury. Furthermore, the extent of myocardial damage of human ischemic heart was accurately assessed by the two signatures of the single-cell and spatial transcriptomic data, suggesting that these two signatures identified from animal model experiments work pretty well for the evaluation of human cardiac function and damage at single-cell and spatial transcriptome levels.

While bulk tissue RNA-seq studies have provided important insights into disease mechanisms contributing to IRI on tissue level, cell type-specific information is lost, and much remains to be learned regarding the roles of individual cell types. The molecular heterogeneity of the different cardiac cell types and changes is evident and known to play crucial roles in human cardiomyopathies including myocardial infarction and heart failure with the use of single-cell technologies (*19, 22, 37, 50-52*). However, whether human cardiac cells of AMI patients received timely PCI therapy are also heterogeneous remained unclear owing to the limited access to the patient heart samples. A recent study elucidated the spatiotemporal heterogeneity of cellular phenotypes in mouse heart with MI/R (*53*). However, the mechanism of time-dependent coronary reperfusion in response to myocardial IRI, particularly metabolic remodeling of cardiac cells, remains unclear. In this study, an appropriate animal model of rat MI/R with different reperfusion time points was used to investigate the time effect of coronary reperfusion on cellular landscape via snRNA-seq. Five major cardiac cell types (CM, FB, EC, MAC, and SMC) were identified from the aforementioned four groups. The CMs were classified into three subclusters based on their characteristics: “healthy”, “subhealthy” and “injured”. The CM1 and CM2 populations were mainly derived from Sham and ETR, respectively. CM3 was attributed to the LTR and MI24h groups. ETR-enriched CM2 exhibited increased lipid oxidation, TCA, and decreased glycolysis but increased heart function signature. Contrarily, LTR and MI24h-enriched CM3 had decreased lipid oxidation, TCA and increased glycolysis but significantly increased heart disease signature. Thus, CMs represent distinct metabolic remodeling states, which critically regulate the heart functions of the CMs within the time-dependent coronary reperfusion followed infarction. The pronounced compositional and transcriptomic alterations in FB, EC and MAC were further analyzed in ETR and LTR. The ETR- and LTR-enriched FB2 induced lipid metabolism accompanied by glycolysis, which further stimulated FB migration and inflammatory response. MI24h-enriched FB3 had high expressions of glycolysis- and ECM-related genes, suggesting that FBs have similar cellular heterogeneity and function in ETR and LTR. ETR had a high proportion of EC2, showing high lipid transport and lipid oxidation scores, and further induced high angiogenesis score. Contrarily, LTR had a high proportion of EC3, which exhibited high expression of glycolytic genes. Our results indicated that reperfusion at different time points (ETR and LTR) significantly affected the metabolic states of cardiac cells, eventually causing distinct cellular function in response to myocardial IRI. ETR maintained energy homeostasis alleviating myocardial IRI by upholding a suitable level of fatty acid oxidation, particularly in CMs.

MACs are broadly divided into M1 and M2 subclusters according to their in vitro construction. Different origins of MACs have been recognized to play distinct roles in heart physiology and diseases, and metabolic states affect MAC function (*54-57*). A previous study demonstrated that snRNA-seq dataset contained a significantly larger number of resident MACs, and the cardiac resident MACs prevented fibrosis and stimulated angiogenesis (*19, 58*). Our study showed that the pro-inflammatory MAC2 and CM2 acted as a hub in cell-cell interactions. The alteration of cell-cell interactions between MACs and CMs was consistent with the studies on normal, failed and partially recovered adult human hearts (*17-19*). Increased Tgfb1 indicated that the anti-inflammatory MAC3 contributed to fibrosis and favored repair of infarcted myocardial tissue in ETR and LTR. The ligand-receptor pairs Ncam1-Cacna1c and Nrg2-Erbb2 increased in ETR than in LTR. Cacna1c and Erbb2 were identified in protecting heart from injury by maintaining calcium homeostasis and activating myocardial protection-related pathways. Furthermore, prediction of the receptors’ downstream target genes indicated that the fatty acid metabolism-related genes *Ppara*, *Ppargc1a*, *Ppargc1b,* and *Ppard* were highly expressed in the “receiver” CM1 and CM2, but not in CM3, which were predicted by the presence of Nampt, Apoe, Igf1 and Bmp6 in the “sender” MACs. Consistently, Nampt has been reported to critically regulate the NAD+ and ATP contents in the heart. The cardiac-specific overexpression of Nampt protects against ischemia-reperfusion injury (*59*). These findings suggested a key cell-cell communication mechanism for ETR maintaining energy homeostasis and decreasing myocardial IRI. The MAC-CM interactions suggested that energy metabolism homeostasis was beneficial not only for promoting anti-inflammatory MACs, but also for protecting myocardial function via cell-cell interaction to maintain the energy of CMs and calcium homeostasis in ETR.

ETR and LTR treatment showed significantly different effects on the energy metabolism remodeling during myocardial IRI. Several lines of evidence helped us to identify Ppara as a core factor for regulating energy metabolism remodeling. In bulk RNA-seq, the expression levels of *Ppara*, *Ppargc1a,* and *Cpt1b* were higher in ETR than in LTR. Simultaneously, in metabolomics analysis, ACar accumulation indicated fatty acid oxidation blockage in LTR. Furthermore, in snRNA-seq, CMs analysis directly revealed that the PPARα motif was significantly enriched in the promoters of energy metabolism-related genes enriched in ETR and significantly downregulated in LTR. PPARα also plays a central role in TF-target gene regulatory module associated with ETR-enriched CM1 and CM2. The PPARA signature, defined by PPARα and its target gene expression, descripted and discriminated the CM metabolic and functional states very well in the analysis of ETR and LTR, or even in the analysis of the single-cell and spatial transcriptomic data of human ischemic heart. RNA velocity analysis from the aspect of cell transiting revealed that the CM2 from the ETR group has a potential to transit to the healthy state. Within the unique CMs in the transiting state, PPARα target genes *Ryr2*, *Cacna1c* and *Myom2* exhibited high dynamic expressions, which were important drivers for the directional flow from CM2 to CM1, suggesting that the inferred directionality of CM2 to CM1 was mainly governed by PPARα and its target genes. Altogether, though those comprehensive analyses, nuclear receptor PPARα was identified as a key regulator for fatty acid oxidation and calcium ion homeostasis in the ETR treatment for myocardial IRI, which enabled us to test the protective effect of PPARα activation in response to myocardial IRI. Fenofibrate serves as an important treatment option for the management of lipid metabolism in patients with diabetes at risk for cardiovascular disease (*60, 61*). Our findings indicated that pretreatment with fenofibrate substantially improved the transcriptional signatures in the rats after IRI. Fenofibrate also significantly upregulated genes associated with heart contraction, fatty acid oxidation and TCA pathways. The IRI induced genes in cardiac cells were downregulated after fenofibrate treatment. A recent study has shown that PPARα pivotally regulates the metabolism, activation, and heterogeneity of MACs and lesion development (*62*). which is consistent with our findings indicating that MAC-enriched genes *S100a8* and *S100a9* were also decreased after fenofibrate treatment. Importantly, angiogenesis-related genes (*Mcam*, *Pdgfa,* and *Bmp2*) were also decreased after this treatment, particularly in ETR, which is consistent with the findings of a previous study indicating that fenofibrate rescues diabetes-related impairment of ischemia-mediated angiogenesis (*63*). Researchers have found that fenofibrate can exhibited anti-inflammatory effects beyond its lipid-lowering (triglycerides in particular) and uric acid-lowering properties (*64, 65*). These results indicated that PPARα pathway activation via fenofibrate treatment improved the transcriptional signatures and function of the IRI hearts in vivo, which was consistent with aforementioned observations in single-nucleus database and explored potential drug repurposing. Overall, our study provides not only a wealth of reference information on cell types and interaction networks in the MI heart with reperfusion at different time points but also insights into the cellular foundation of cardiac metabolism homeostasis and the potential that PPARα activation could serve as a therapeutic target for IRI.

## Methods

### Rat model of myocardial ischemia/reperfusion

Rat underwent in situ myocardial ischemia and reperfusion injury as described previously (*66, 67*). The rats were anesthetized by 1-3% isoflurane and connected to a ventilator (Comerio, ITALY) via tracheal intubation. After exposing the heart, the left anterior descending coronary artery was ligated for 1h or 6h with 7/0 nylon suture and thereafter released for a 23h or 18h period for reperfusion. Sham operated rats received the same surgery without ligation. The chest was closed after the surgery had been completed, and warmed till recovery.

### Fenofibrate injection

Rats were treated by intraperitoneal injection with fenofibrate (100 mg/kg) 2h before the LAD surgery, and PBS was used for the control.

### Echocardiogram

Transthoracic echocardiography was performed using the Type B Ultrasonography system (Vinno 6, China). Rats were anaesthetized by 2% isoflurane inhalation and placed on a heated platform (37°C). Heart rate was stabilized between 450-500 beats per minute before echocardiographic measurements. Left ventricle and the aortic outflow tract were observed under the two-dimensional B-Mode, and the sample line was placed at the maximum cross-section of left ventricle to guide the recording of serial M-Mode echocardiographic images. The fraction shortening (FS), ejection fraction (EF), left ventricular end-diastolic internal diameter (LVIDd) and left ventricular end-systolic internal diameter (LVIDs) were measured from at least three distinct frames for each rat.

### Histological studies

After MI/R and MI operations, the heart samples of Sham, MI1h, MI6h, MI1h/R, MI6h/R and MI24h groups were harvested and rinsed three times in PBS and arrested in 10% KCl solution. Tissues were processed routinely for histology and embedded in paraffin, sectioned at 4 microns thickness stained with H&E, Masson trichrome staining and Terminal deoxynucleotidyl transferase dUTP nick end labeling (TUNEL) assay according to the manufacturers.

### RNA isolation and qRT-PCR

Total RNA was extracted following the manufacturer’s instructions using the RNeasy RNA isolation Kit (Qiagen) and reverse transcribed to cDNA with the HiScript III 1st Strand cDNA Synthesis Kit (Vazyme). The Real-time PCR reactions were performed with Taq Pro Universal SYBR qPCR Master Mix (Vazyme) using ABI Prism 7500 system. The primer sequences were documented in the supplementary table.

### Bulk sequencing and data analysis

RNA-seq was performed at Novogene Corporation. In brief, total RNA was purified from the infarct zone of Sham, MI and MI/R using RNeasy RNA isolation Kit (Qiagen) coupled with an on-column DNase I digestion to eliminate residual DNA. The mRNA was purified from 2 µg of total RNA, and libraries for massive parallel sequencing were constructed with NEBNext® UltraTM RNA Library Prep Kit for Illumina (NEB). Library quality was assessed with Agilent Bioanalyzer 2100 system. Next-generation sequencing was performed on an Illumina HiSeq 4000 platform. Paired-end 150 bp reads of bulk RNA-seq data were aligned to rn7 reference genome (ENSEMBL, version 67) using STAR alignment tool (version 2.7.3a) with default parameters (*68*). Uniquely mapped reads were employed to quantify gene expression using RSEM software (version 1.3.1) (*69*). Transcripts per million mapped reads (TPM) were calculated to estimate the gene expression levels, normalized for sequencing depth. Euclidean distance clustering and principal component analysis (PCA) based on TPM values were used to evaluate the similarities and variabilities across biological replicates and groups. Differential gene expression (DGE) analysis was performed using DESeq2 R package, those genes with adjusted P-value < 0.05 and absolute fold change > 2 were assigned to differentially expressed genes (DEGs). Unbiased clustering of variable genes across different time points were performed using Mfuzz package. Those genes with clustering membership > 0.6 were retained in clusters and visualized using pheatmap R package.

### Single-nucleus isolation and single-nucleus RNA-sequencing

Nuclei were isolated with Nuclei EZ Lysis buffer (NUC-101; Sigma-Aldrich) supplemented with protease inhibitor (5892791001; Roche) and RNase inhibitor (N2615; Promega and AM2696; Life Technologies). Heart samples were cut into 2-mm pieces and homogenized using a Dounce homogenizer (885302-0002; Kimble Chase) in 2 ml of ice-cold Nuclei EZ Lysis buffer, and they were incubated on ice for 5 minutes with an additional 2 ml of lysis buffer. The homogenate was filtered through a 40-mm cell strainer (43-50040-51; pluriSelect) and then centrifuged at 5003g for 5 minutes at 4°C. The pellet was resuspended and washed with 4 ml of the buffer, and then, it was incubated on ice for 5 minutes. After another centrifugation, the pellet was resuspended in Nuclei Suspension Buffer (1xPBS, 0.07% BSA, and 0.1% RNase inhibitor), filtered through a 20-mm cell strainer (43-50020-50; pluriSelect), and counted. Collected nuclei for each sample were loaded onto 10X Chrominum platform. cDNA libraries were constructed using Chrominum Single-Cell 3’GEM, Library & Gel Bead Kit v3, then sequenced on illumina novaseq 6000.

### The snRNA-seq data processing

The reads were aligned to rat reference genome (rn7; ENSEMBL, version 67) using CellRanger v6.1.1 with default parameters, excluding the option ‘-include-introns’ to obtain intronic reads in nuclei. Only uniquely mapped reads without duplications were retained. This provided a unique molecular identifier (UMI) counting matrix of 48,637 nuclei with 32,623 genes. In the filtering process, we firstly filtered cells with less 1000 UMI counts or more than 20,000 UMI counts. Those nuclei expressing less than 5,00 genes with a minimum of 5% mitochondrial genes were also discarded. Genes expressing in a minimum of 20 cells were retained. Finally, a gene matrix of 22,234 nuclei and 17,003 genes were used for downstream analysis using Seurat (v4.0.2) under R (4.0.5) environment. Briefly, standard Seurat “integration” pipeline was performed to remove batch effect across different samples. Then the raw UMI counts were normalized to the total reads with log-normalization and scaled by a factor of 10,000, top 2,000 highly variable genes selected by Variance Stabilizing Transformation (VST) were used to perform principal component analysis (PCA). Utilizing 1st to 30th principal components (PCs), dimension reduction was implemented by Uniform Manifold Approximation and Projection (UMAP). Further cell clusters were identified with 1st to 30th PCs based on the Louvain-clustering. And differentially expressed genes (DEGs) in each cluster were identified using “FindAllMarkers” function in Seurat with default parameters.

### Clusters annotation

Cluster annotation were performed using ‘SingleR’ R package based on the annotation from previous publication (*17*). Then canonical marker genes were used to verify the annotated cell-types. Briefly, the cluster 0 detected expression of *Dcn* and *Pdgfra* was assigned to fibroblasts (FBs). The cluster 1 highly expressing *Actn2* and *Myh6* was annotated as cardiomyocytes (CMs). Two clusters, cluster 2 and cluster 5, were assigned to endothelial cells (ECs) based on the expression of *Vwf* and *Kdr*. Cluster 3 and cluster 6, highly expressing *Ptprc* (*CD45*) and *Cd68*, were labeled as macrophages (MACs). Cluster 5 highly expressing *Mylk* and *Rgs5* was annotated as smooth muscle cells (SMCs).

### Cell trajectory analysis

Pseudo-time trajectory analysis was performed using the R package Monocle2 with standard inputs and default parameters. Top 2,000 variable genes across CMs sub-clusters were employed to perform dimension reduction of “DDRTree”. After ordering cells, a tree-like cell trajectory was constructed. Python package Velocyte was used to estimate the RNA velocity in each major cell type, visualization was presented by scVelo.

### Transcription factor regulon analysis

We calculated the Pearson correlation between single nuclei of DEGs across three CM sub-clusters. For each sub-cluster, those genes with correlation coefficient > 0.3 or < −0.3 were retained. Based on the annotation of AnimalTFDB (http://bioinfo.life.hust.edu.cn/AnimalTFDB4), highly variable transcription factor genes were obtained from DEGs of CM sub-clusters and regarded as potential regulons. Regulation networks for highly variable transcription factors and correlated genes were constructed by the GENIE3 R package, then visualized by Cytoscape.

Motif position weight matrix (PWM) files of 119 rat transcription factors were downloaded from MEME website (https://meme-suite.org/meme). Promoter was defined as the 2,000 bp window around transcription start sites (TSSs) of genes. Using R packages ‘Biostrings’ and ‘BSgenome.Rnorvegicus.UCSC.rn7’, we extracted the promoter sequences of CM1-high DEGs, CM3-high DEGs and non-DEGs respectively. Then scanned 119 TF motifs around promoter sequences of above gene sets. Using non-DEGs as background, further Fisher’ exact test was performed to identify the over-represented TF motifs in CM1-high DEGs and CM3-high DEGs respectively. Adjusted *P*-value < 0.05 and odds ratio > 1 were used as statistical significances to identify enriched TF motifs. The sequence logos of enriched motifs were plotted in by seqLogo package.

### Cell-cell interaction analysis

Utilizing LIANA (v0.1.10), we inferred significant ligand–receptor interactions (*P*-value < 0.01) among the previously generated sub-cluster Seurat objects, including CMs, FBs, ECs and MACs (https://github.com/saezlab/liana). Next, we used CrossTalker (v1.3.5) to find changes in cell–cell communication by contrasting ligand– receptor interactions predicted in MI1h/R vs MI6h/R and MI1h/R vs MI24h/R respectively. NicheNet (v1.1.1) was implemented to predict potential targets in receiver cell according to detailed instructions (https://github.com/saeyslab/nichenetr).

### Gene pathway analysis

Gene Ontology (GO) enrichment analysis of differentially expressed genes was implemented by the clusterProfiler R package. The *p*-values were adjusted using the Benjamini-Hochberg method. GO terms with adjusted *p*-value less than 0.05 were considered significantly enriched by differential expressed genes.

### Metabolomics data generation and analysis

Metabolomic data analyses were conducted at Metabolon as described previously using nontargeted metabolomic profiling (*70*). Heart samples from Sham, MI1h/R, MI6h/R and MI24h groups were subjected to methanol extraction and then were split into aliquots for analysis by ultrahigh performance liquid chromatography/mass spectrometry (UHPLC/MS) in the positive, negative or polar ion mode and by gas chromatography/mass spectrometry (GC/MS). Metabolites were identified by automated comparison of ion features to a reference library of chemical standards followed by visual inspection for quality control. For statistical analyses and data display, any missing values were assumed to be below the limits of detection; these values were imputed with the compound minimum (minimum value imputation). To determine statistical significance, significance analysis of microarrays (SAM) was conducted on the residuals from a multiple regression model, which included the age and sex as covariates. The adjusted *P*-value <0.05 was used as an indication of high confidence in a result. A total of 67 differentially regulated metabolites were observed in IML versus IMH individuals. Pathway analysis was conducted using MetPA24, which combines several advanced pathway enrichment analyses along with the analysis of pathway topological characteristics across more than 874 metabolic pathways. For over-representation and pathway topology analyses, a hypergeometric test and the relative-betweenness centrality were used, respectively.

### Immunostaining

Immunostaining was performed according to standard protocols for paraffin embedded samples. For paraffin sections, hearts were collected in PBS on ice in 4% PFA at room temperature for 12-24h depending on tissue size, followed by tissue dehydration and embedding. Embedded hearts samples were sectioned to a thickness of 4 μm. Prior to staining, sections were dewaxed, and antigen was retrieved with citric acid solution, followed by incubation with BSA, and the primary antibody was incubated overnight at 4 °C or 1-3h at room temperature. The next day, tissue sections were washed 3 times with PBS, incubated with Alexa-Fluor-conjugated secondary antibodies for 1.5 h at room temperature, washed 3 times with PBS, and mounted with mounting medium containing the nuclear stain DAPI. The following antibodies were used (ratios indicate antibody dilution): TNNT2 (Abcam, ab8295; 1:500), TNNI3 (Abclonal, A0152; 1:250), NPPA (Abclonal, A22075; 1:250), PGC1α (Proteintech, 66369-1-Ig; 1:250). Alexa Fluor 488 AffiniPure Goat Anti-Rabbit IgG (H+L) Alexa Fluor (Yeasen, 33106ES60; 1:500) and Dylight 594, Goat Anti-Mouse IgG (Abbkine, A23410; 1:500) were used for secondary antibody incubation. Fluorescent images were captured using SLIDEVIEW VS200 Research Slide Scanner (Olympus, Japan).

### Western blotting

Fresh heart tissue was lysed in high-salt buffer (20 mM HEPES pH 7.4, 500 mM NaCl, 25% glycerol, 0.2 mM EDTA, 0.5 mM DTT, 1% NP40 and 0.5% SDS plus protease) on ice for 15 min. The lysate was centrifuged at 12000 rpm for 10min, and the supernatant was boiled with 5x Loading buffer. The extracted protein was separated in SDS-PAGE gel, transferred to 0.22 um PVDF membrane and probed by the subsequent incubation with primary antibodies and HRP-conjugated secondary antibodies. The target proteins were detected with the enhanced chemiluminescent substrate. ImageJ was employed for expression quantification.

### Data accessibility

The snRNA-seq data and bulk tissue RNA-seq have been submitted to NCBI’s Gene Expression Omnibus (GEO) and are accessible through GEO accession numbers GSE240850.

### Statistical analysis

All results are expressed as means ± SEM. Student’s t-test or Mann-Whitney Wilcoxon rank-sum test was used for comparison of 2 groups as indicated, and one-way analysis of variance (ANOVA), then by post hoc analysis was used for comparison of 3 or more groups as indicated in the manuscript. P value < 0.05 was considered statistically significant. The *p*-values in multiple tests were adjusted to false discovery rate (FDR) using the Benjamini-Hochberg method. All statistical analyses were performed using R software (version 4.0.3).

## Acknowledgements

This study is financially supported by the National Key Research and Development Program of China (2021YFA0805902, 2019YFE0117400, 2022YFA1103204) and the National Natural Science Foundation of China (32270884, 82202328, 81974025, 32022023). The authors thank research staff Jinrong Peng from College of animal sciences of Zhejiang university for his assistance on the animal studies, and Jun Chen from College of life sciences of Zhejiang university for his assistance on the histological analysis of rat heart.

## Author contributions

L.Z. performed conceptualization and supervision. L.S., Y.Z. and G.S. performed experimental design and execution. Y.G., F.H., and F.R. performed single-cell RNA-seq analysis. D.L. and J.H. performed bulk RNA-seq analysis. Y.G. and Z.Z. performed metabolic analysis. J.Z., L.S. and Y.L. performed rat MI/R model. Y.G., Y.Z. G.S. performed echocardiography and Histomorphometry analysis. F.H., F.R., and Y.L. collected heart tissues, preformed western blot, conducted and analyzed qPCR experiments. Z.L. Y.G. L.S. and Y.Z. prepared the manuscript. All authors reviewed and approved the manuscript. The authors declare no competing interests.

## Supplementary Information for

**Fig. S1.**
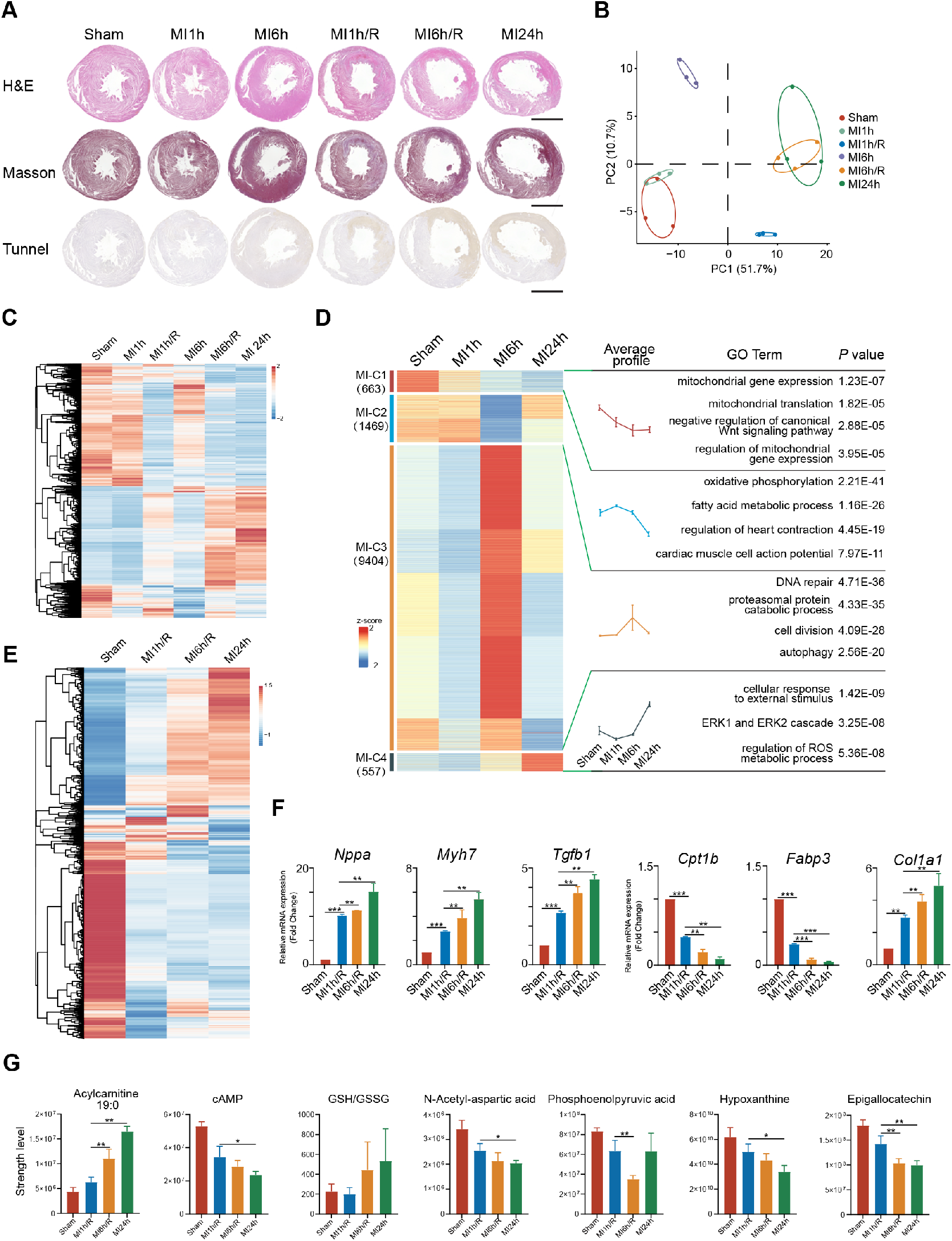
Early time reperfusion reduced myocardial IRI by maintaining fatty acid metabolism homeostasis. **(A)** Histological analysis of Sham, MI and MI/R heart (up: H&E staining; middle: Masson trichrome staining; down: Tunnel assay). **(B)** PCA analysis based on the Sham. MI and MI/R groups. **(C)** Heatmap of DEGs identified from group Sham and MI24h in the six groups (Sham, MI and MI/R). **(D)** Heatmap of DEGs identified from group Sham and MI24h in the four groups (Sham, MI1h/R, MI6h/R and MI24h). **(E)** Average expression levels and the Gene Ontology (GO) analysis of each clustered genes were shown by the heatmap and temporal profile of Sham, MI1h, MI6h and MI24h groups. **(F)** The expression levels of key genes were measured by *qRT-PCR*. **(G)** Bar graphs showing relative abundance of the crucial metabolites.

**Fig. S2.**
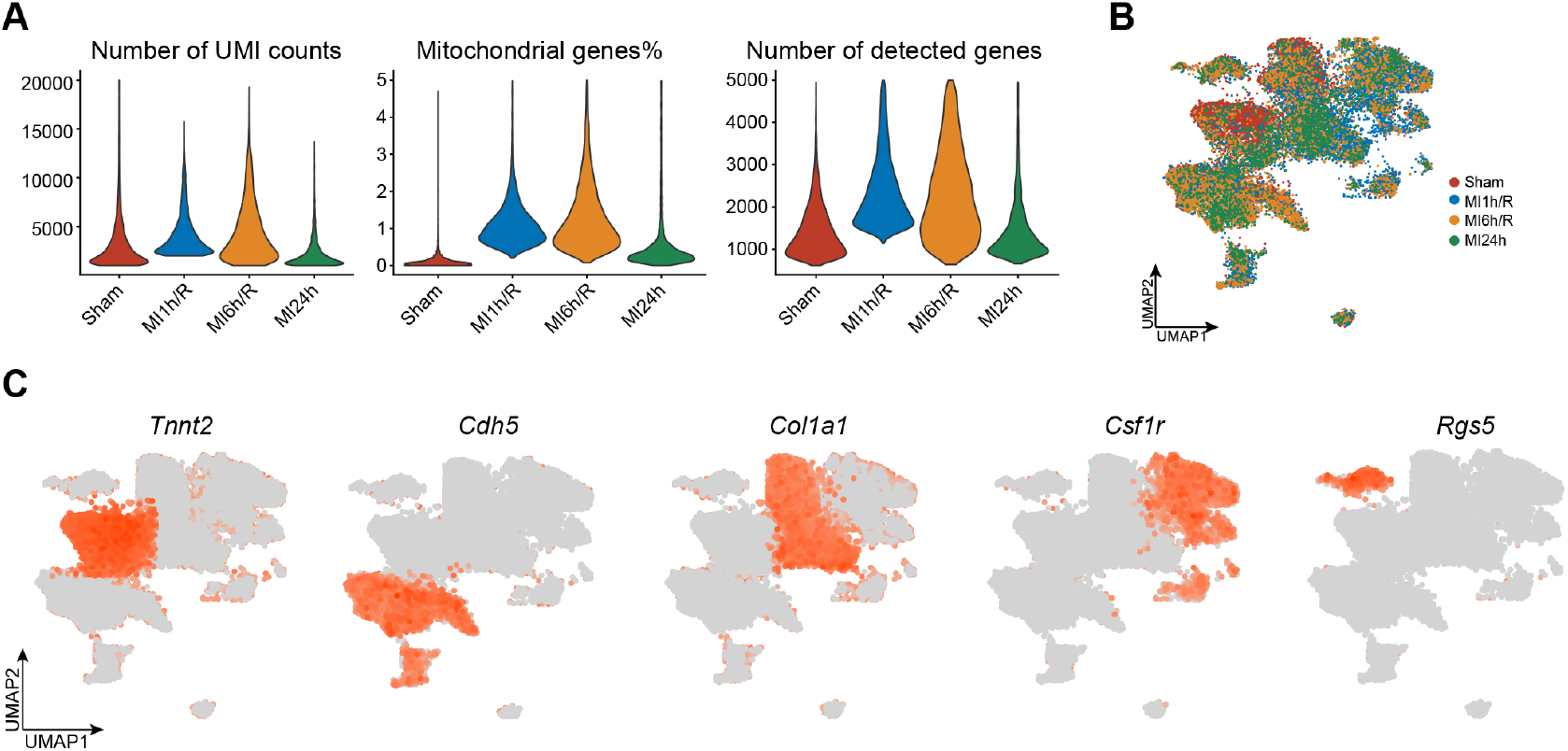
Single-nucleus transcriptomes revealed the cellular landscape of the cell composition of the IRI hearts. **(A)** Quality metrics for single-nucleus RNA-Seq data showing number of UMI counts in the four groups (left), proportion of mitochondrial genes (middle), and number of genes detected per group (right). **(B)** UMAP plot of 22,234 single cells isolated from four groups. Cells are colored by group. **(C)** Feature plots showing expression level of cell markers of CM, EC, FB, MP and SMC.

**Fig. S3.**
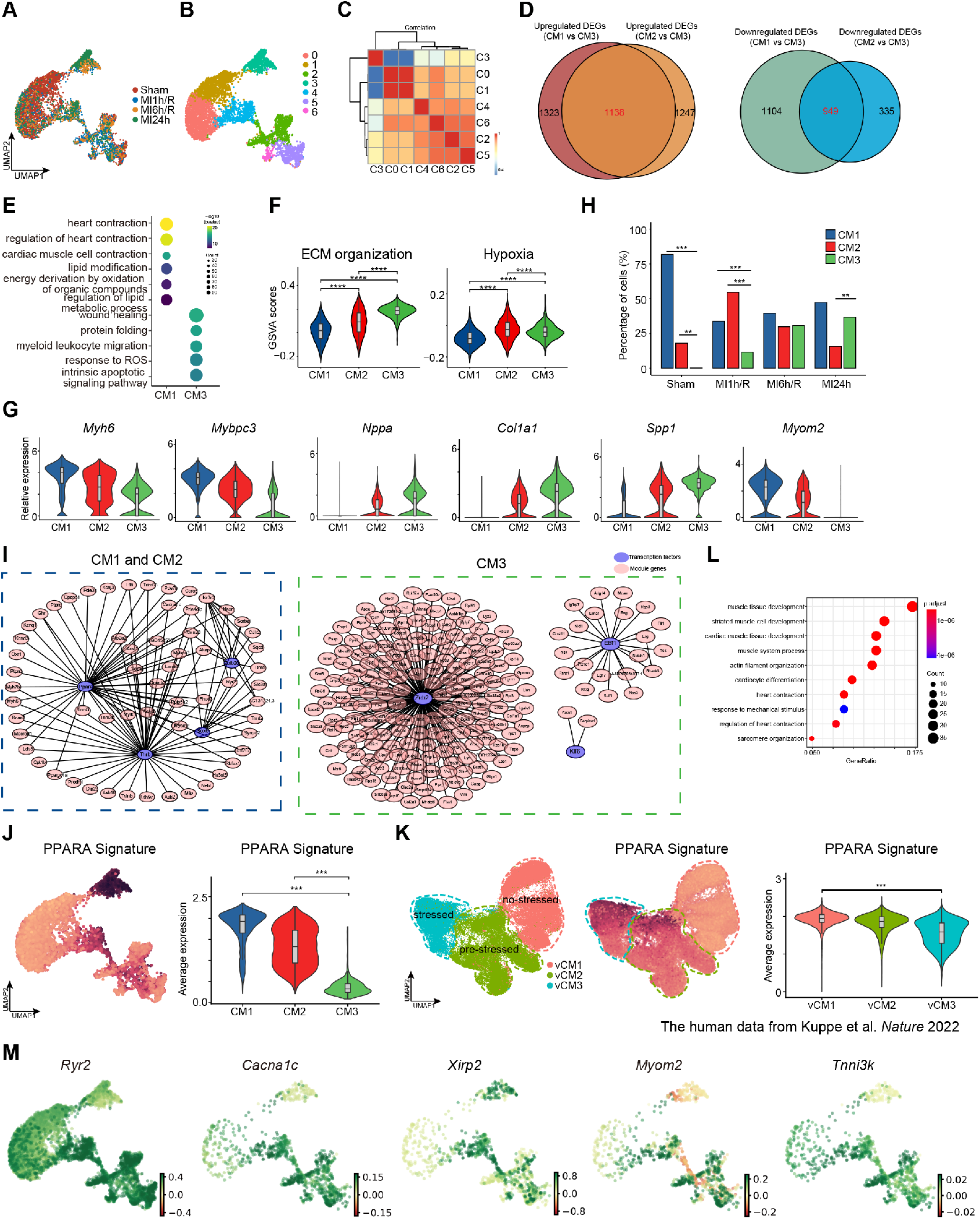
Early time reperfusion safeguarded PPAR*α*-mediated fatty acid metabolism in cardiomyocytes. **(A-B)** UMAP projection showing 5,239 CMs isolated from rat hearts at different times of reperfusion. Cells are color labelled by group **(A)** and subclusters **(B)**. **(C)** The correlation among the 7 CM subclusters. **(D)** Venn diagram shows the overlap of upregulated (The left panel) and downregulated (The right panel) genes between CM1,2 and CM3. **(E)** GO enrichment for the DEGs identified from CM1 and CM3. **(F)** GSVA analysis of the important signaling pathways. **(G)** Violin plots showing the expression level of crucial genes in 3 CM subclusters. **(H)** Statistical analysis of subclusters CM populations based on the groups. **(I)** TF target network created from CM1 and CM2-High genes (The left panel), CM3-High genes (The right panel). **(J)** Feature plot and violin plots showing the expression level of PPARA signature in the 3 CM subclusters. **(K)** UMAP projection showing CMs in the published human myocardial infarction heart single-cell sequencing datasets (The left panel), feature plot (The middle panel) and violin plots (The right panel) showing the expression level of PPARA signature in 3 the human CMs. **(L)** GO enrichment for the DEGs identified from transiting CM2. **(M)** Expression Level of putative driver genes showing the extra variation drived by the transiting.

**Fig. S4.**
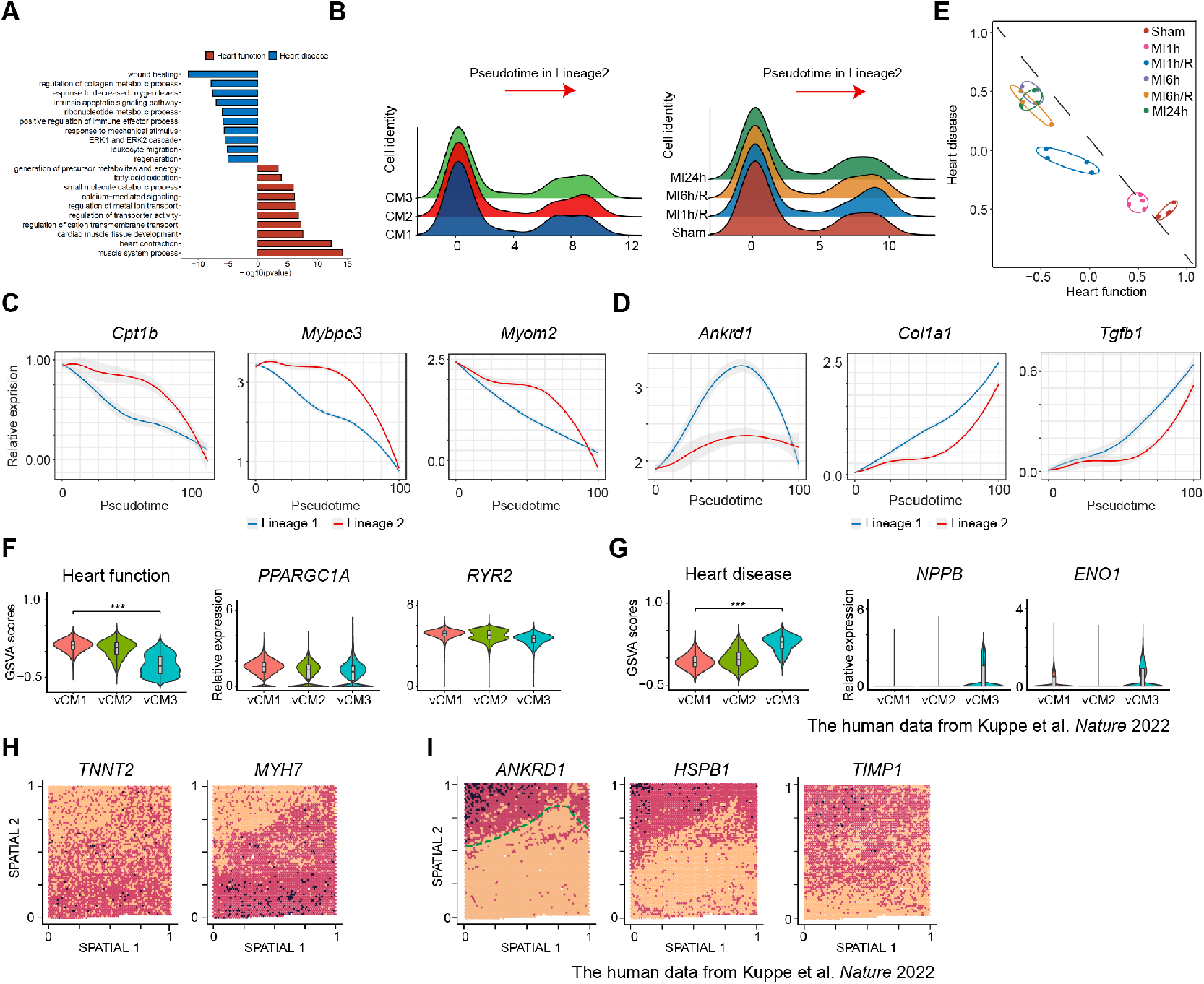
Heart function and heart disease signatures were identified to assess the myocardial injury. **(A)** The GO enrichment analysis for genes of heart function and heart disease signatures. **(B)** Density plots for cell density of the 3 CM subclusters (The left panel) and the four groups (The right panel) along with the pseudo-time lineage 2. **(C)** Expression for the representative heart function genes over pseudo-time. **(D)** Expression for the representative heart disease genes over pseudo-time. **(E)** The analysis of heart function and heart disease signatures based on the bulk tissues transcriptomics datasets of the six groups. **(F)** Violin plots showing the expression of heart function signatures and representative genes in the CMs from published human heart single-cell sequencing datasets. **(G)** Violin plots showing the expression of heart disease signatures and representative genes in the CMs from published human heart single-cell sequencing datasets. **(H-I)** Violin plots showing the expression for representative genes of heart function **(H)** and heart disease **(I)** signature from published human heart single-cell sequencing datasets.

**Fig. S5.**
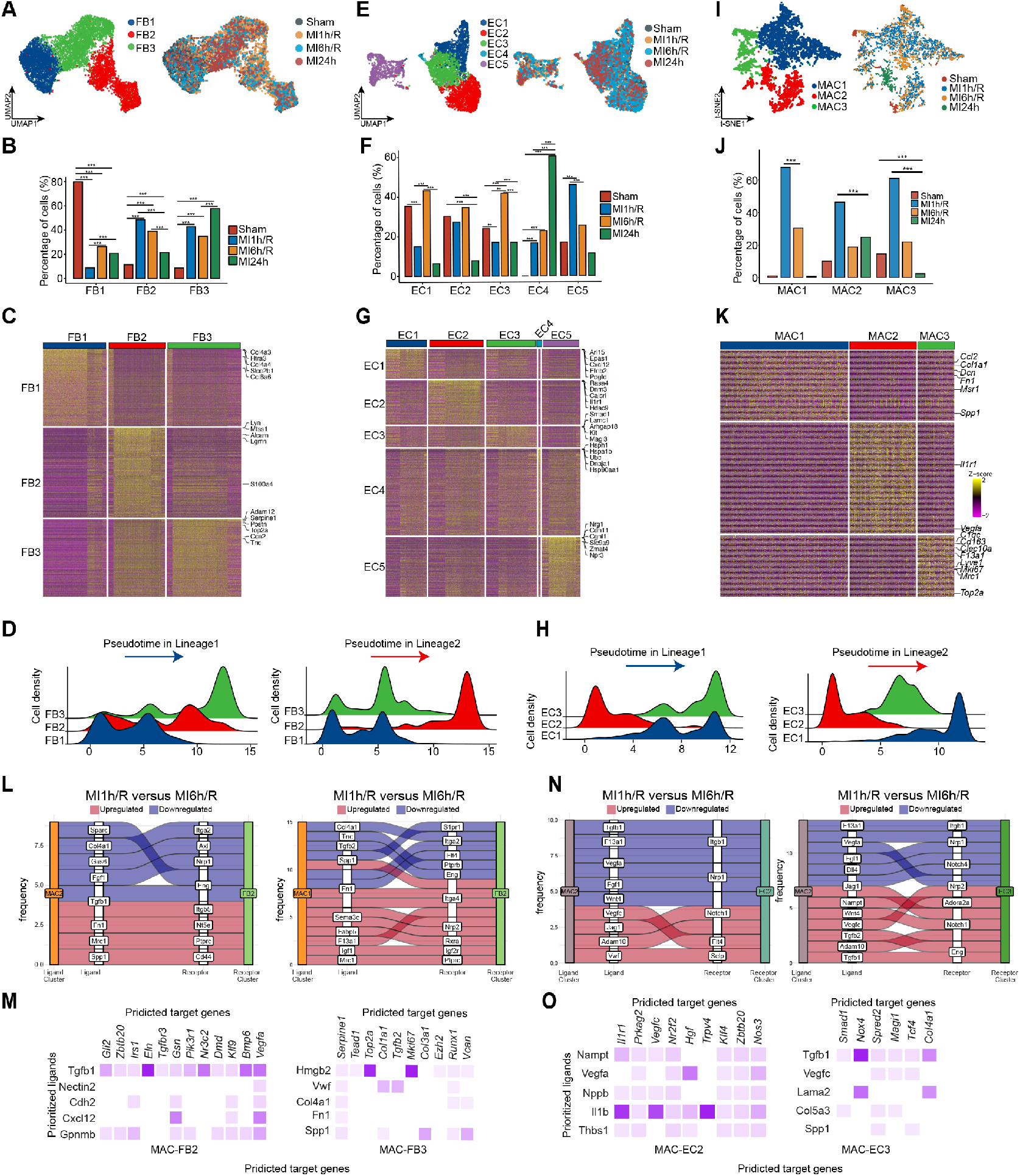
The metabolic remodeling affects the function of cardiac FBs and ECs as well as the cell-cell interactions of macrophages contributed to remodeling of cardiac cells. **(A)** UMAP projection showing FBs isolated from rat hearts at different times of reperfusion. Cells are color labelled by subclusters (the left panel) and groups (the right panel). **(B)** Statistical analysis of the subclusters FB populations. **(C)** Heatmap showing enriched genes identified from FB1-3. **(D)** Density plots for cell density of the 3 FB subclusters with the pseudo-time lineage 1 and 2. **(E)** UMAP projection showing ECs isolated from rat hearts at different times of reperfusion. Cells are color labelled subclusters (the left panel) and groups (the right panel). **(F)** Statistical analysis of the subclusters FB populations based on the group. **(G)** Heat map showing enriched genes identified from EC1-5. **(H)** Density plots for cell density of the 3 EC subclusters with the pseudo-time lineage 1 and 2. **(I)** The tSNE projection showing MAC isolated from rat hearts at different times of reperfusion. Cells are color labelled by subclusters (the left panel) and groups (the right panel). **(J)** Statistical analysis of the subclusters MAC populations. **(K)** Heat map showing differentially expressed genes catalogued in the DEGs identified from the 3 clusters. **(L)** Sankey plots of ligand-receptor interactions for MAC1/ MAC2 and FB2 identified from MI1h/R versus MI1h/R. These ligand-receptor pairs are selected by absolute value of the difference in LRscore. **(M)** The heatmap of select ligands expressed by MAC-FB2/3 to regulate the genes which are differentially expressed by the FB2/3. Well-established ligand-target gene interactions shown with a darker shade of purple. **(N)** Sankey plots of ligand-receptor interactions for MAC2 and EC2/3 identified from MI1h/R versus MI1h/R. These ligand-receptor pairs are selected by absolute value of the difference in LRscore. **(O)** The heatmap of select ligands expressed by MAC-EC2/3 to regulate the genes which are differentially expressed by the EC2/3. Well-established ligand-target gene interactions shown with a darker shade of purple.

**Extended Data Fig.6.**
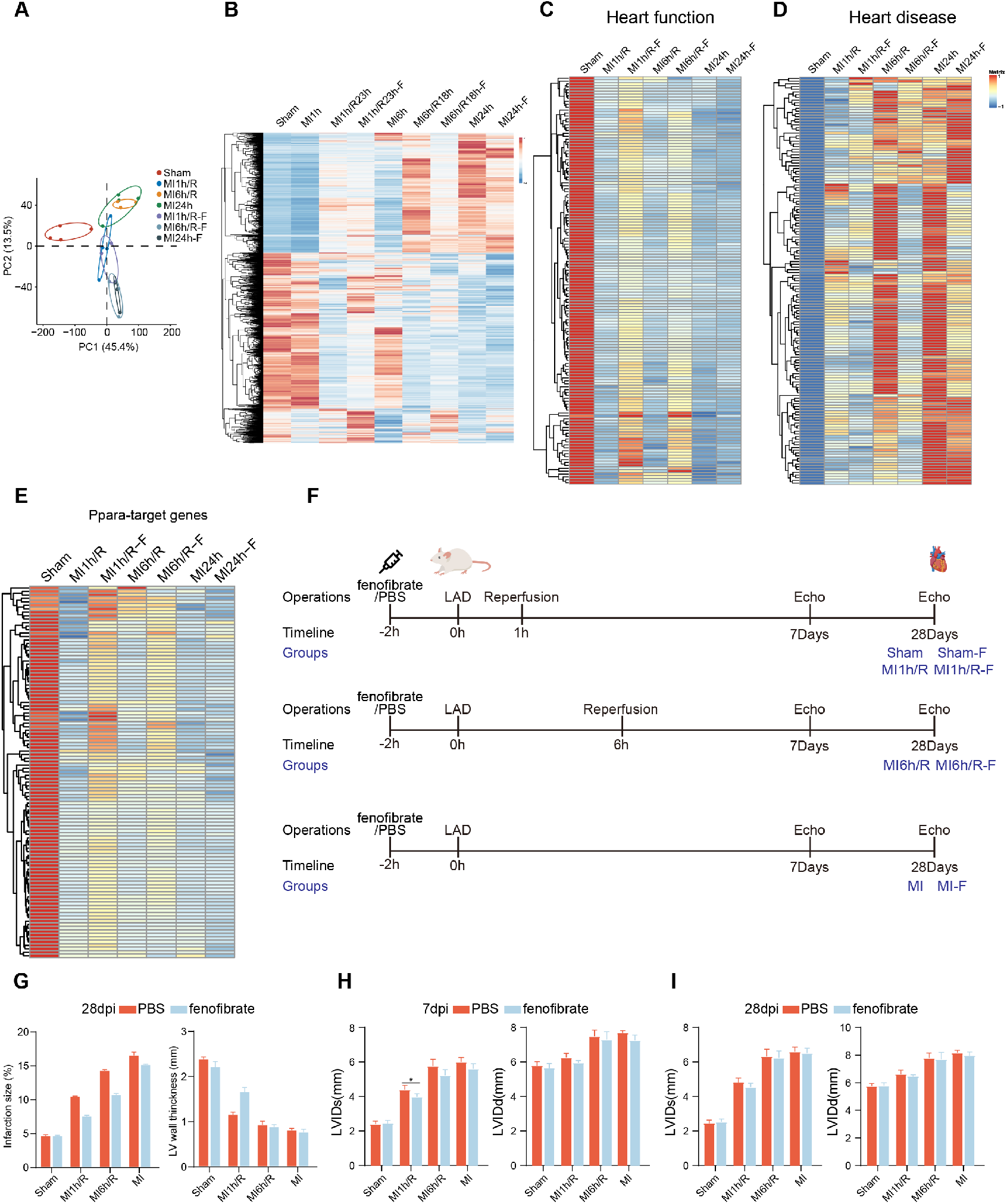
Pretreatment with fenofibrate ameliorated myocardial IRI. **(A)** PCA analysis by bulk tissues transcriptomics datasets with PPARα agonist fenofibrate. **(B)** Heatmap showed the expression of DEGs with and without fenofibrate. **(C-D)** Heatmap showed the expression of heart function **(c)** and heart disease genes **(d)** with and without fenofibrate. **(E)** Heatmap showed the expression of Ppara-target genes with and without fenofibrate. processes. **(F)** Schematic of study design and the rats were treated with PPARα agonist fenofibrate 2 hours before LAD, and ECHO was performed 7days and 28days after MI. **(G)** Morphometric parameters including the percentage of infarction size and wall thickness in MI region were measured via Image J software. **(H)** Each bar plot represents the left ventricle diastolic internal diameter (LVIDd) and left ventricle systolic internal diameter (LVIDs) systolic at day 7 days post infarction. **(I)** Each bar plot represents the left ventricle diastolic internal diameter (LVIDd) and left ventricle systolic internal diameter (LVIDs) systolic at day 28 days post infarction.

